# Long Noncoding RNAs Preserve Pancreatic Cancer Identity and Resist Cell Fate Conversion

**DOI:** 10.1101/2025.06.25.661419

**Authors:** Dmytro Grygoryev, Seung-Won Lee, Connor Mitchell Frankston, Shauna Rakshe, Mark Berry, Alex Hirano, Taelor Ekstrom, Elise Manalo, Julien Tessier, Marilynn Chow-Castro, Jason Link, Dove Keith, Brett C. Sheppard, Suzanne Fei, Terry Morgan, Helen E. Remotti, Wenli Yang, Emma E. Furth, Sudhir Thakurela, Rosalie Sears, Jungsun Kim

**Affiliations:** Cancer Early Detection Advanced Research Center, Knight Cancer Institute, Oregon Health & Science (OHSU) School of Medicine, Portland, OR; Department of Molecular and Medical Genetics, OHSU School of Medicine, Portland, OR; Biomedical engineering graduate program, Department of Biomedical Engineering, OHSU School of Medicine, Portland, OR; Biostatistics Shared Resource, Knight Cancer Institute, OHSU, Portland, OR; Lurie Center for Autism, Department of Pediatrics, Massachusetts General Hospital, Harvard Medical School, Boston, MA; Brenden-Colson Center for Pancreatic Care, OHSU School of Medicine, Portland, OR; Department of Surgery, OHSU School of Medicine, Portland, OR; Knight Cancer Institute, OHSU School of Medicine, Portland, OR; Pathology & Laboratory Medicine, OHSU School of Medicine, Portland, OR; Department of Pathology, Columbia University, New York, NY; Institute for Regenerative Medicine, UPenn Perelman School of Medicine, Philadelphia, PA; Pathology and Laboratory Medicine, UPenn Perelman School of Medicine, Philadelphia, PA; Center for Genomic Medicine, Massachusetts General Hospital, Boston, MA; Broad Institute of MIT and Harvard, Cambridge, MA, USA

**Keywords:** cancer cell identity, transcriptional memory, transcriptional rigidity, long noncoding RNA, master transcription factor-mediated cellular reprogramming, pancreatic cancer, pluripotency, plasticity, *ATXN7L3-AS1*, *AC079921.2*

## Abstract

The Yamanaka factors (OCT4, SOX2, KLF4, and MYC; OSKM) can rejuvenate aging phenotypes in somatic cell types by resetting the epigenetic landscape. Curiously, most solid tumor cells remain largely resistant to reprogramming despite their well-documented plasticity, and the underlying mechanisms are unclear. Here, we combined genomic profiling and *in vivo* assays to investigate OSKM-mediated reprogramming of pancreatic ductal adenocarcinoma (PDAC). In the initial stages, we found that cancer-specific genes were refractory while mesodermal/ECM programs, normally silenced by PRC2, were aberrantly upregulated. A CRISPR interference screen for OSKM reprogramming coupled with functional analyses revealed that suppression of cancer-associated long noncoding RNAs (lncRNAs) erased malignant epithelial programs, restored tumor suppressor activity, and impaired tumorigenicity *in vivo*. We further identified that *ATXN7L3-AS1* lncRNA sustains the PDAC malignant identity through its association with active epithelial oncogenic programs and poised PRC2-targeted developmental genes, thereby supporting both plasticity and memory. Thus, by exploring why cancer cells are resistant to reprogramming, we identify lncRNAs as gatekeepers of malignant identity, suggesting that targeting lncRNAs could be a generalizable therapeutic strategy in treating solid tumors.

## Introduction

Cellular reprogramming through forced expression of lineage-defining transcription factors (TFs), including pioneer factors, can erase established cell identity and impose new cell fates^1–7^. The first demonstration of induced pluripotency was achieved by combined expression of the TFs OCT4, SOX2, KLF4, and MYC (OSKM, or the Yamanaka factors), which reprogrammed normal somatic cells into induced pluripotent stem cells (iPSCs)^6, 7^. Beyond generating iPSCs, OSKM expression has been reported to rejuvenate aging phenotypes by reversing age-associated epigenetic alterations^8, 9^. Because many cancers segregate with aging and are partly driven by epigenetic dysregulation, reprogramming has been proposed as a potential strategy for resetting the malignant cell fate.

However, with the exception of neuronal lineage and hematologic malignancies^10–12^, solid tumors exhibit broad resistance to OSKM reprogramming^13–17^. While combined expression of the four factors partially reprograms solid cancer cells, the resulting states are often unstable: cells either stall in intermediate states^13^ or revert to malignant phenotypes^14, 15^. Persistent ectopic factor expression can attenuate malignant features and maintain a “tamed” state; yet these cells remain refractory and do not undergo complete reprogramming^16^. In some contexts, ectopic expressions of OSKM or partial reprogramming even reinforce tumor-like programs^17^. These findings suggest that reprogrammed cancer cells retain a transcriptional and epigenetic memory of their tissue of origin and therefore resist complete erasure of cell identity^18, 19^. Given that the plasticity of cancer cells is well-documented ^20–25^, this striking resistance suggests the existence of robust and cancer-specific transcriptional memory mechanisms. Yet, the molecular basis of such a memory mechanism remains poorly understood.

PDAC, one of the deadliest human cancers in aging patients^26^, exemplifies the paradox of a cell state that is both plastic and fixed. In PDAC cancer cells, transcriptional transitions between classic and basal subtypes are dynamic^27–31^. Yet, cells maintain a stable malignant identity defined by oncogenic programs and ductal epithelial signatures that underpin therapeutic resistance^32, 33^. Identifying mechanisms that can maintain this malignant identity, despite its dynamic transcriptional states, is crucial for developing effective therapeutic strategies against this lethal cancer.

Gene regulatory networks comprise TFs, cofactors, and chromatin regulators that tightly coordinate the maintenance of cell identity^34–37^. Recently, long noncoding RNAs (lncRNAs) have emerged as key epigenetic regulators, exhibiting greater tissue specificity than most protein-coding genes^38–41^. By scaffolding chromatin-remodeling complexes and guiding gene expression in trans and cis, lncRNAs are well poised to orchestrate cell-type-specific transcriptional programs^42–49^. As such, they have essential roles in embryonic development, pluripotency, and differentiation^49–53^, and influence reprogramming by modulating microRNAs that control core pluripotency factors^53, 54^. Conversely, dysregulation of lncRNAs is increasingly recognized as a disease-driving mechanism in cancer, contributing to both disease progression and therapeutic resistance^55–58^. Yet, specific lncRNAs in solid tumors such as PDAC that act to preserve malignant identity and block cell fate conversion have not been identified.

Here, we leveraged PDAC as an aging-associated cancer model to investigate how OSKM reprogramming is constrained in the context of cancer-specific transcriptional memory. Specifically, we coupled a systematic CRISPR interference (CRISPRi) screen with OSKM reprogramming, integrative genomic analyses, and *in vivo* xenograft assays. Through this approach, we identified a subset of cancer-associated lncRNAs that enforce the coexistence of plasticity and epithelial memory, thereby preserving malignant ductal identity and impeding reprogramming.

## RESULTS

### PDAC Cells Resist OSKM-Reprogramming, Revealing Restricted Plasticity

PDAC comprises two major transcriptional subtypes: classic and basal-like (also referred to as squamous)^27–29^. In prior work, we implanted human primary PDAC tissue into immunodeficient mice to establish and fully characterize patient-derived xenograft (PDXs) tumor models, thereby avoiding selection against PDAC during reprogramming (Table S1)^59^. We thus performed OSKM reprogramming in PDAC cells representing both classical (CFPAC1, ST-00014490 PDX) and basal subtypes (Panc1, ST-00007599 PDX) ^59, 60^ using BJ fibroblasts and H6c7 human normal pancreatic duct epithelial cells (hnPDECs) as normal controls (Fig. 1A).

**Figure 1.**
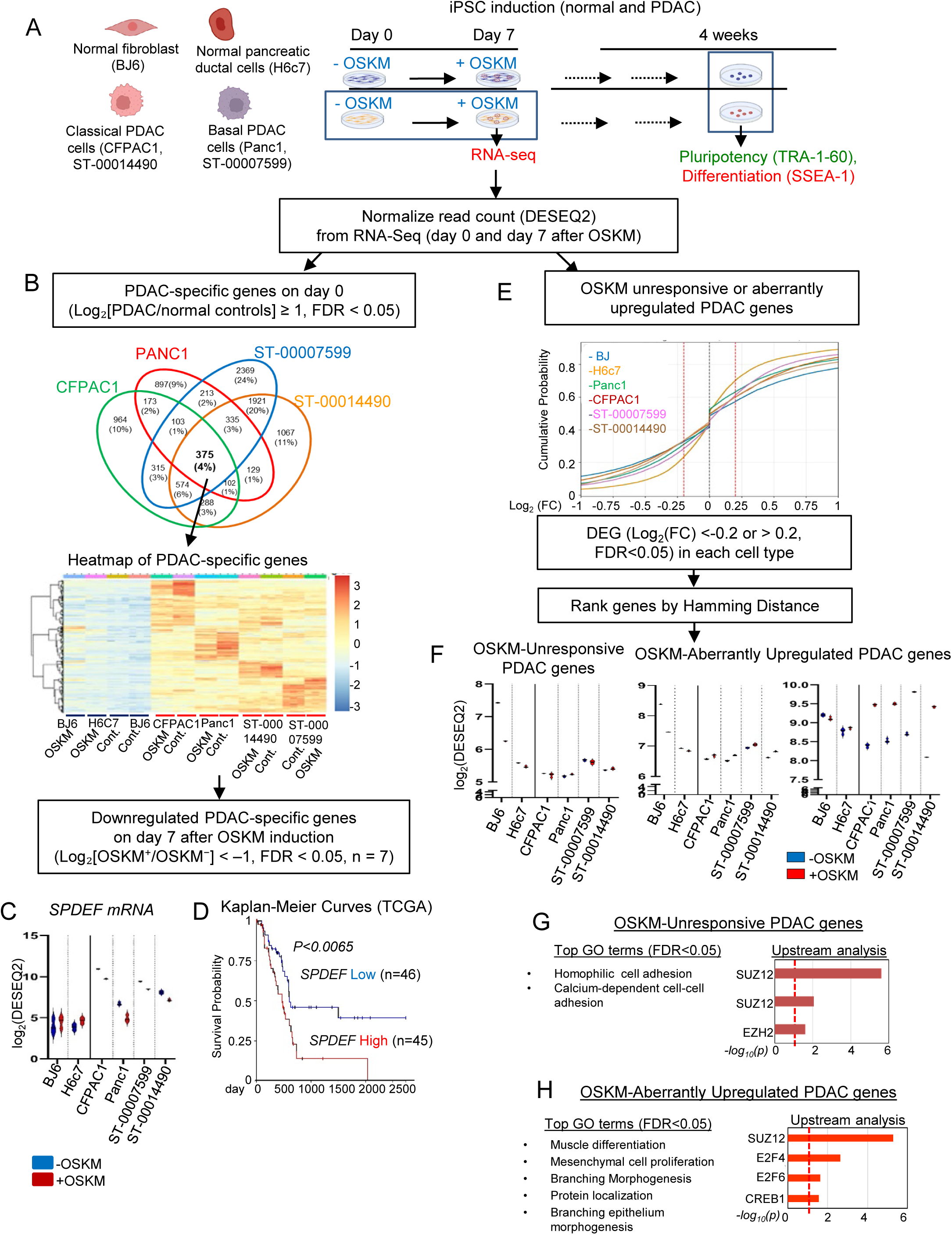
OSKM Reprogramming Partially Suppresses PDAC-Unique Genes, Leaving Others Unresponsive or Aberrantly Induced. A. Schematic diagram of reprogramming of normal controls (BJ fibroblast and H6c7 hnPDEC) and human classical (CFPAC1, ST-00014490) and basal (Panc1, ST-00007599) subtypes of PDAC cells using Sendai virus-mediated expression of OSKM factors. Cells were transduced with OSKM-expressing Sendai virus on day 0. For transcriptomic analysis, cells were harvested on days 0 and 7. To assess pluripotency and early differentiation, cells were fixed on day 30 and immunostained for TRA-1-60 and SSEA1, respectively. B. Each Venn diagram shows PDAC-specific genes that are upregulated relative to normal controls on day 0 (log₂ [PDAC/normal controls] ≥ 1, FDR < 0.05). The heatmap displays the expression profiles of 375 PDAC-specific genes consistently upregulated across all PDAC samples, with Z-scores indicated by the color key. Seven of the 375 PDAC-unique genes were significantly downregulated by OSKM across all PDAC cells (log₂[OSKM⁺/OSKM⁻] < –1, FDR < 0.05). C-D. Example of an OSKM-induced downregulated PDAC-specific gene. (C) *SPDEF* mRNA levels in the indicated cells without and with OSKM induction. (D) Kaplan–Meier survival analysis of *SPDEF* expression in the TCGA PDAC cohort (n = 150). Patients were stratified by SPDEF expression quartiles (Low: <7.611; High: >9.675), based on log₂-transformed normalized RSEM values. P-value was calculated using the log-rank test (χ², rho = 0). The high *SPDEF* expression in PDAC is associated with poor overall survival of patients with PDAC. E. Identification of aberrantly increased or unchanged genes in PDAC following OSKM induction. The empirical cumulative distribution function (ECDF) displays the cumulative probability distribution of log₂ fold-change (FC) values across samples, before and after OSKM induction (day 7 vs. day 0). The X-axis indicates log₂(FC), and the Y-axis indicates the cumulative fraction of genes. The plot is zoomed in to the range of −1 to 1 to highlight subtle shifts in gene expression. Genes with log₂(FC) > 0.2 or < –0.2 and FDR < 0.05 were defined as DEGs in each sample, based on inspection of the Mean Absolute Deviation (MAD) heatmap as an empirical measure of deviation from the null hypothesis (Fig. S3C; see Methods). Genes were then queried against pre-defined patterns of expression changes across the samples. A weighted Hamming distance was used to measure how well a gene’s pattern of expression changes across samples matched with each query (see Methods). For each query, genes with weighted Hamming distances no greater than 0.35 were selected as hits for downstream analysis. F. Violin plots showing expression levels for OSKM-unresponsive PDAC genes (left: distance < 0.375, n = 1,466; Table S2F) and OSKM-aberrantly upregulated genes (middle: distance < 0.25, n = 1,815; right: distance = 0, n = 82; Tables S2G–S2H; see Methods). G-H. Gene Ontology (GO)^102^ and X2K upstream regulator^103, 104^ analyses for genes that were either unresponsive to OSKM in PDAC (G, Table S2F) or aberrantly upregulated following OSKM induction (H, Table S2G). The red dashed line indicates the significance threshold (FDR or p-value < 0.05).

Instead of transgene-mediated reprogramming, we utilized Sendai virus to deliver the OSKM-expressing polycistronic vector^60^ as it provides a clean platform for reprogramming somatic cells without causing genomic integration^61^. Importantly, this system achieved efficient and consistent transgene expression across all primary PDAC cells, comparable to that in H6c7 hnPDECs and BJ fibroblasts^60^.

We first validated the Sendai virus–mediated OSKM delivery system using BJ fibroblasts and acinar cells isolated from pancreatitis tissue of PDAC patients (Fig. S1A-S1B). The reprogramming efficiency of acinar cells (0.4%) was comparable to that of fibroblasts (Fig. S1C). The resulting iPSC lines expressed the canonical pluripotency marker TRA-1-60^62, 63^, exhibited normal karyotypes, and successfully differentiated into all three germ layers (Fig. S1D-F), validating that Sendai virus–mediated OSKM delivery effectively reprogrammed human pancreatic acinar cells and fibroblasts into a pluripotent state.

In contrast to pancreatic acinar cells, H6c7 hnPDECs failed to express TRA-1-60 by day 30 post-OSKM induction. However, many cells expressed SSEA-1, a marker associated with pluripotency in mouse cells and with early embryonic differentiation in human cells^64, 65^ (Fig. S2A). Because TRA-1-60 expression emerged by day 45-60^66^, we concluded that hnPDECs retain a stronger lineage memory than acinar cells or fibroblasts. When we conducted OSKM induction in classical-type PDAC cells, we detected expression of SSEA-1 but not TRA-1-60, consistent with stalled progression toward pluripotency (Fig. S2A). Similarly, in OSKM-induced basal subtype PDAC cells, TRA-1-60 expression was sporadic while SSEA-1 expression was unaffected, indicating resistance to transcriptional rewiring (Fig.S2A). Even after 60 days of OSKM induction, PDAC cells failed to achieve full reprogramming. Altogether, these findings indicate that PDAC cells maintain stable, cancer-associated programs that prevent complete cell fate conversion.

### OSKM Overrides Only a Subset of PDAC Identity Genes

Early molecular events following OSKM induction are critical in determining cell fate outcomes^3, 67–73^. To characterize such early transcriptional responses, we examined transcriptomic profiles of PDAC cells (ST-00007599, ST-00014490, Panc1, CFPAC1) and control cells (BJ fibroblasts and H6c7 hnPDECs) at day 7 post-OSKM induction (Fig. 1A). Notably, ectopic OSKM expression levels were comparable across all cell types (Fig. S2B), with Principal Component Analysis (PCA) showing consistent patterns across technical replicates within each sample (Fig. S3A).

OSKM initiates reprogramming of somatic cells by silencing their original lineage-specific gene programs^3, 67–73^. We therefore hypothesized that OSKM might downregulate a subset of the PDAC-specific transcriptome, particularly those involved in the malignant identity. To test this hypothesis, we defined PDAC-unique genes by identifying transcripts significantly upregulated across all PDAC cells relative to both fibroblasts and hnPDECs prior to OSKM reprogramming (Fig. 1B, Table S2A). Out of 375 PDAC-unique genes, we only found 7 that were significantly downregulated on day 7 following OSKM induction (Fig. 1B, Table S2B), several of which were associated with apoptosis and epithelial differentiation pathways (Table S2C). These findings suggest that early reprogramming in PDAC is constrained by restrictive regulatory mechanisms. Notably, one of the OSKM-responsive genes was SPDEF, which encodes the ETS transcription factor with a SAM-pointed domain^74^, a newly reported driver of classical PDAC tumorigenesis^75^. As expected, we found that high expression of SPDEF correlated with poor prognosis (Fig. 1C-D).

Next, we assessed transcriptional responses to OSKM 7 days after induction across different cell types. We detected the greatest number of differentially expressed genes (DEGs) in BJ fibroblasts. In contrast, we detected fewer in H6c7 hnPDECs, and markedly fewer DEGs in PDAC cells overall (Fig. S3B, Table S2D). These variations likely reflect inter-sample heterogeneity, as we detected more DEGs in each PDAC sample than in H6c7 hnPDECs (Fig. S3B, Table S2E) but also indicate higher intrinsic transcriptional plasticity in PDAC. Importantly, this plasticity was not accompanied by loss of malignant identity. Instead, it was constrained by robust transcriptional memory that preserved PDAC-specific programs despite OSKM induction.

Thus, while OSKM partially suppressed tumor-associated gene programs, the core transcriptional identity of PDAC cells remained largely intact.

### PRC2-Linked Lineage Programs Persist During Reprogramming

Based on our data thus far, we hypothesized that OSKM-unresponsive genes in PDAC act as gatekeepers that preserve malignant identity, whereas OSKM-aberrantly upregulated genes (silenced or unchanged in normal cells) represent maladaptive, cancer-specific barriers that antagonize cell fate conversion. Such aberrant responses could potentially drive maladaptive PDAC plasticity as well as impeding reprogramming.

To identify such mechanisms, and account for inter-sample heterogeneity, we developed a classification framework incorporating variability across PDAC (n=4) and normal (n=2) samples. DEGs before and after OSKM induction were identified using likelihood ratio testing (LRT)^76^ and ranked by a weighted Hamming distance^77^ relative to predefined patterns. These included: (1) downregulated in normal cells but minimally changed in cancer (“OSKM unresponsive in PDAC”), and (2) unchanged or downregulated in normal cells but upregulated in cancer (“OSKM aberrantly upregulated in PDAC”) (Fig. 1E-F and S3C–D, Table S2F-H, method). While OSKM-unresponsive genes were enriched in cell adhesion pathways, OSKM-aberrantly upregulated genes were associated with mesodermal and morphogenetic lineage programs (Fig. 1G–H). Both categories were predicted targets of the PRC2 core components (SUZ12, EZH2) (Fig. 1G-H) and significantly overlapped with known SUZ12 targets in human and mouse embryonic stem cells (mES and huES) (Fig. S3E). Notably, expression of the PRC2 subunit remained unchanged post-OSKM (Fig. S3F), suggesting that impaired PRC2 recruitment—rather than reduced subunit abundance—underlies the failure to suppress these programs.

In summary, OSKM downregulated a small subset of PDAC marker genes and left most PDAC-specific genes unchanged or upregulated. Many of these belonged to PRC2-regulated lineage programs.

### Suppressing Cancer-Associated lncRNAs Unlocks Reprogramming Efficiency

Given the broad role of lncRNAs in regulating epigenetic machinery, including recruiting PRC2^42, 78–82^, we next performed CRISPRi screens targeting 691 cancer-associated lncRNAs (10 gRNAs per lncRNA plus controls; 7,087 gRNAs total; Table S3A) to identify those sustaining malignant identity^83, 84^. The screens were conducted in the context of OSKM-mediated reprogramming of normal ductal cells and PDAC cells, using the H6c7 and HPAFII cells expressing catalytically inactive Cas9 (dCas9)-KRAB (henceforth referred to as H6c7-dCas9-gRNA and HPAFII-dCas9-gRNA; Fig. 2A).

**Figure 2.**
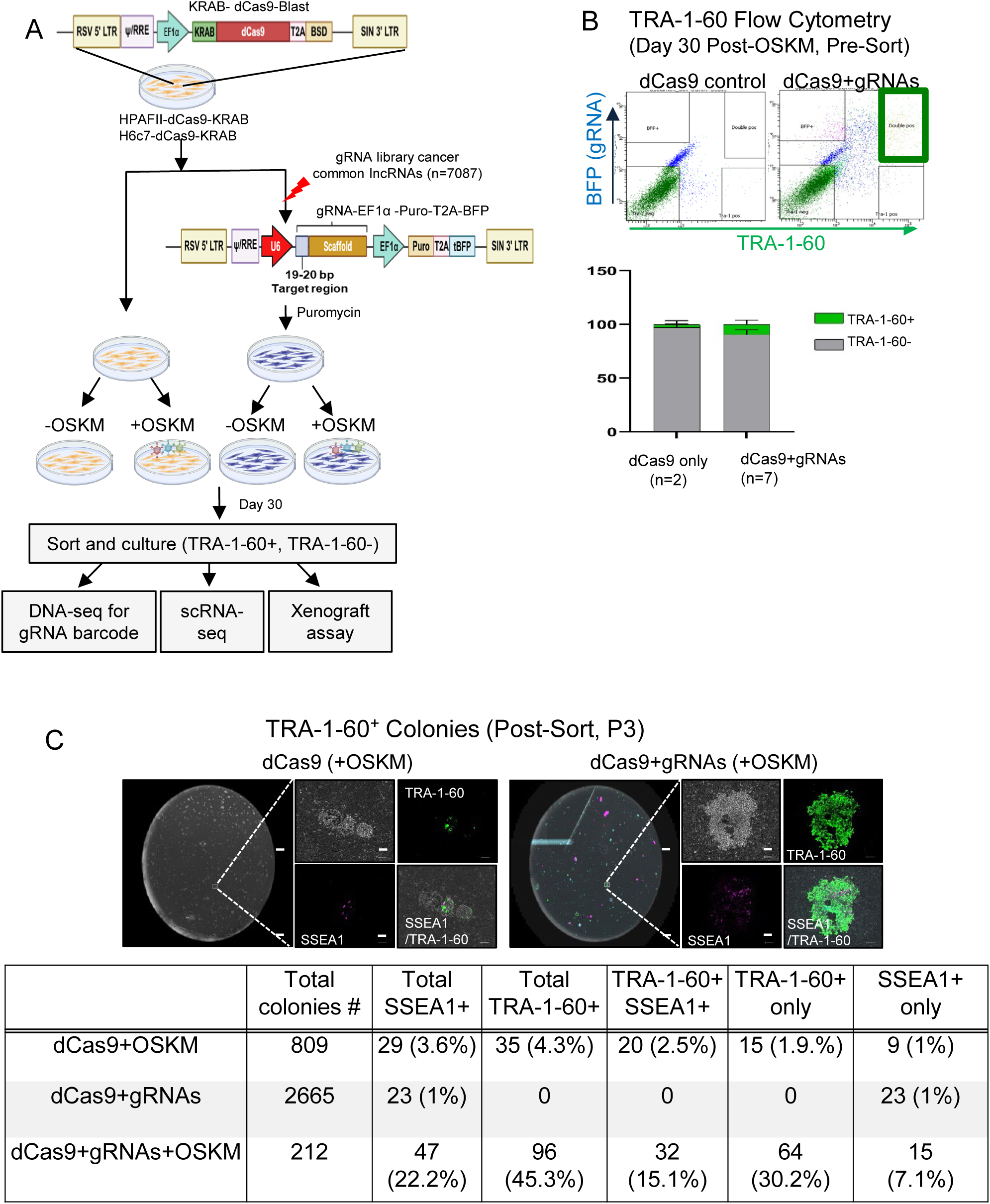
CRISPRi of Cancer-Associated lncRNAs Enhances Reprogramming Efficiency in PDAC Cells. A. Schematic diagram of reprogramming HPAFII PDAC and H6c7 hnPDEC cells by CRISPRi screening. The reprogramming efficiency was measured by immunostaining for TRA-1-60 and SSEA1. The TRA-1-60+ pluripotent stem cell-like cells were quantified and sorted by flow cytometry at 30 days. B. Flow cytometry–based quantification of TRA-1-60⁺ and TRA-1-60⁻ fractions in HPAFII-dCas9 control cells and HPAFII-dCas9 cells expressing lncRNA-targeting gRNAs (BFP⁺) at day 30 after OSKM induction. C. Representative images and quantification summary of TRA-1-60⁺ and SSEA1⁺ cells on plates after multiple passages (Scale bar: 284 μm).

As expected, TRA-1-60⁺ pluripotent-like colonies were rare in HPAFII dCas9 control cells by day 30 post-OSKM induction. In contrast, cells transduced with the lncRNA-targeting gRNA pool showed an increased frequency of TRA-1-60⁺ cells (Fig. 2B). The isolated TRA-1-60⁺ and TRA-1-60⁻ populations (hereafter referred to as dCas9+gRNA+OSKM-T^+^ and dCas9+gRNA+OSKM-T^−^, respectively) remained phenotypically stable over multiple passages (Fig. 2C). In contrast, the same gRNA pool targeting lncRNAs did not enhance reprogramming of H6c7 hnPDEC cells (Fig. S3G), suggesting that these lncRNAs act in a PDAC-specific manner to preserve malignant transcriptional memory.

### Pooled lncRNA Depletion Enables OSKM to Erase PDAC Identity and Restore Tumor Suppression

Next, we profiled single-cell transcriptomes from OSKM-induced, pooled-CRISPRi PDAC cells (dCas9+gRNA+OSKM-TRA-1-60⁺ [T+] and TRA-1-60⁻ [T-]) (Fig. 3A, S4A–B) to assess whether depletion of cancer-associated lncRNAs was sufficient to override PDAC cell identity and establish a new gene program (pluripotency). While Louvain clustering resolved 15 distinct clusters, control dCas9 cells localized mainly to tubular ductal (Group 0) and invasive epithelial (Group 1) clusters, consistent with PDAC identity (Fig. 3B–C, S4C). As expected, OSKM expression alone was insufficient to erase PDAC programs (Fig. 3B-C, S4C). Similarly, dCas9–gRNA cells displayed metabolic shifts (pyruvate metabolism up, oxidative phosphorylation down; Fig. S4D) yet remained confined to PDAC-like clusters (Fig. 3B–C).

**Figure 3.**
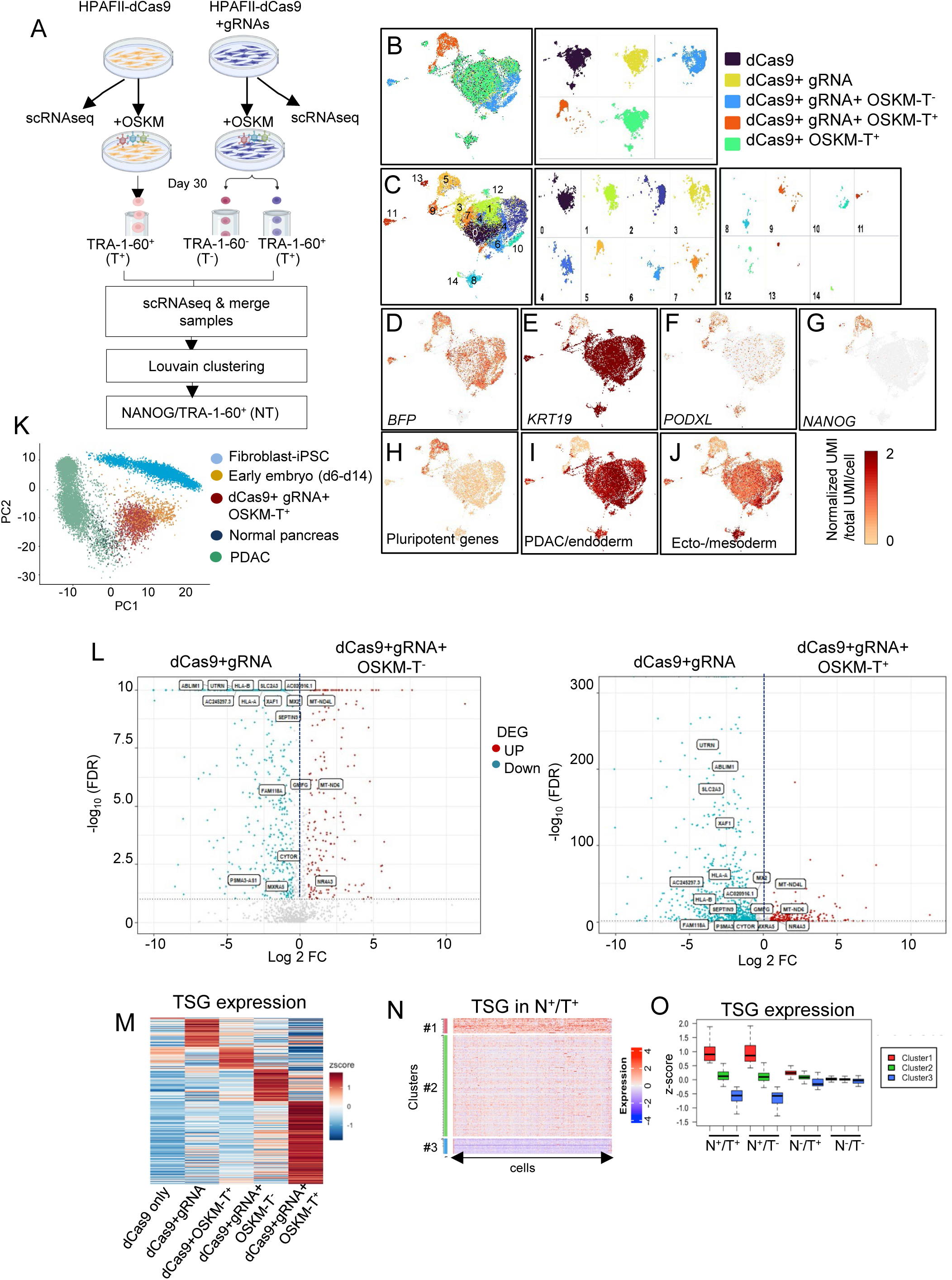
Repression of PDAC Programs and De-repression of Pluripotency and Tumor Suppressor Signatures in PDAC Cells Reprogrammed with OSKM and lncRNA Depletion. A. Schematic diagram of the scRNA-seq workflow. B-J. UMAP visualizations showing distinct clusters by sample (B), 15 Louvain-defined clusters (C), cells expressing BFP (D), pancreatic ductal marker KRT19 (E), *PODXL* encoding TRA-1-60 (F), *NANOG* (G), pluripotency-associated genes (H, Table S4A), endodermal and PDAC-related genes (I, Table S4B), and ectodermal/mesodermal lineage markers (J, Table S4C). Each dot represents a cell, and the color intensity indicates the normalized UMI counts (scaled by the total UMI per cell). K. PCA of HPAFII-dCas9+gRNA+OSKM-T^+^ cells alongside publicly available scRNA-seq datasets from human PDAC (n=6, GSE212966^83^), human and mouse normal pancreas (n=4, GSE84133^84^), human early embryo before the three germ layers formed (D6-D14, n=555, GSE136447^105^), and iPSC derived from Parkinson’s disease patients’ fibroblast (n=12, GSE183248^106^). L. Volcano plots of DEGs between dCas9+gRNA and either dCas9+gRNA+OSKM-T^−^ (left) or dCas9+gRNA+OSKM-T^+^ (right). PDAC-specific signature genes are annotated (n=332, Table S4D). M. Heatmap showing expression of 318 TSGs across the indicated cell populations (Table S4E). The plot shows z-scores of the mean TSG expression across cells in the sample. The distinct subsets show that each sample has a set of TSGs more highly expressed in that sample than in others. N-O. Heatmap (N) and quantification (O) of TSG expression in the dCas9+gRNA+OSKM-T^+^ cells. (N) Each column represents an individual cell, each row corresponds to a TSG, and expression levels are shown as Z-scores normalized across all cells for each gene. TSGs were grouped into three distinct clusters based on their expression profiles using unsupervised k-means clustering. (O) Expression levels of TSGs in each cluster are shown across the indicated subpopulations based on NANOG and TRA-1-60 expressions-NANOG⁻/TRA-1-60⁻ (N⁻/T⁻), NANOG⁻/TRA-1-60⁺ (N⁻/T⁺), NANOG⁺/TRA-1-60⁻ (N⁺/T⁻), and NANOG⁺/TRA-1-60⁺ (N⁺/T⁺). The Y-axis indicates Z-score.

In contrast, dCas9+gRNA+OSKM-T^+^ cells redistributed away from PDAC clusters (Groups 0 and 1) and localized predominantly to Group 5, characterized by pluripotency-associated gene expression and loss of pancreatic/germ layer markers (Fig. 3B–J, S4E-F, Table S4A–C). PCA confirmed that T^+^ cells occupied a transcriptional space distinct from PDAC or normal pancreas and aligned most closely with early human embryo profiles (Fig. 3K).

Both T^+^ and T^−^ populations were depleted of mucus-producing and invasive epithelial clusters (Fig. 3B–C), indicating a broad disruption of malignant programs. A curated PDAC-specific gene signature (Table S4D)^85, 86^ was significantly downregulated only in the combined lncRNA-depleted OSKM condition (Fig. 3L). While controls expressed few tumor suppressor genes (TSGs), dCas9+gRNA+OSKM-T^+^ cells showed robust and broad TSG re-expression, enriched within NANOG⁺ states (Fig. 3M–O), based on comparative analysis of the OncoKB TSG (Table S4E)^87, 88^.

Together, these results demonstrate that pooled suppression of cancer-associated lncRNAs during reprogramming erases PDAC transcriptional identity and produce pluripotent-like states where tumor suppressor programs are reactivated, confirming the reversal of malignant fate.

### Pooled lncRNA Depletion with OSKM Erases PDAC Identity and Limits Tumorigenesis In Vivo

We next investigated xenograft outcomes in immunodeficient NOD-SCID-IL2Rγ^null^ (NSG) mice^89^ to assess whether depleting PDAC identity-associated transcriptional programs would tame cancer progression *in vivo* (Fig. 4A). As expected, parental HPAFII cells formed tumors within one-month after injection, with dCas9-gRNAs producing slightly smaller tumors compared to their parental (Fig. 4B–C). Strikingly, mice injected with dCas9+gRNA+OSKM-T^+^ cells either failed to develop tumors or formed significantly smaller tumors than dCas9+gRNA+OSKM-T^−^ and all controls (p < 0.005; Fig. 4B–C).

**Figure 4.**
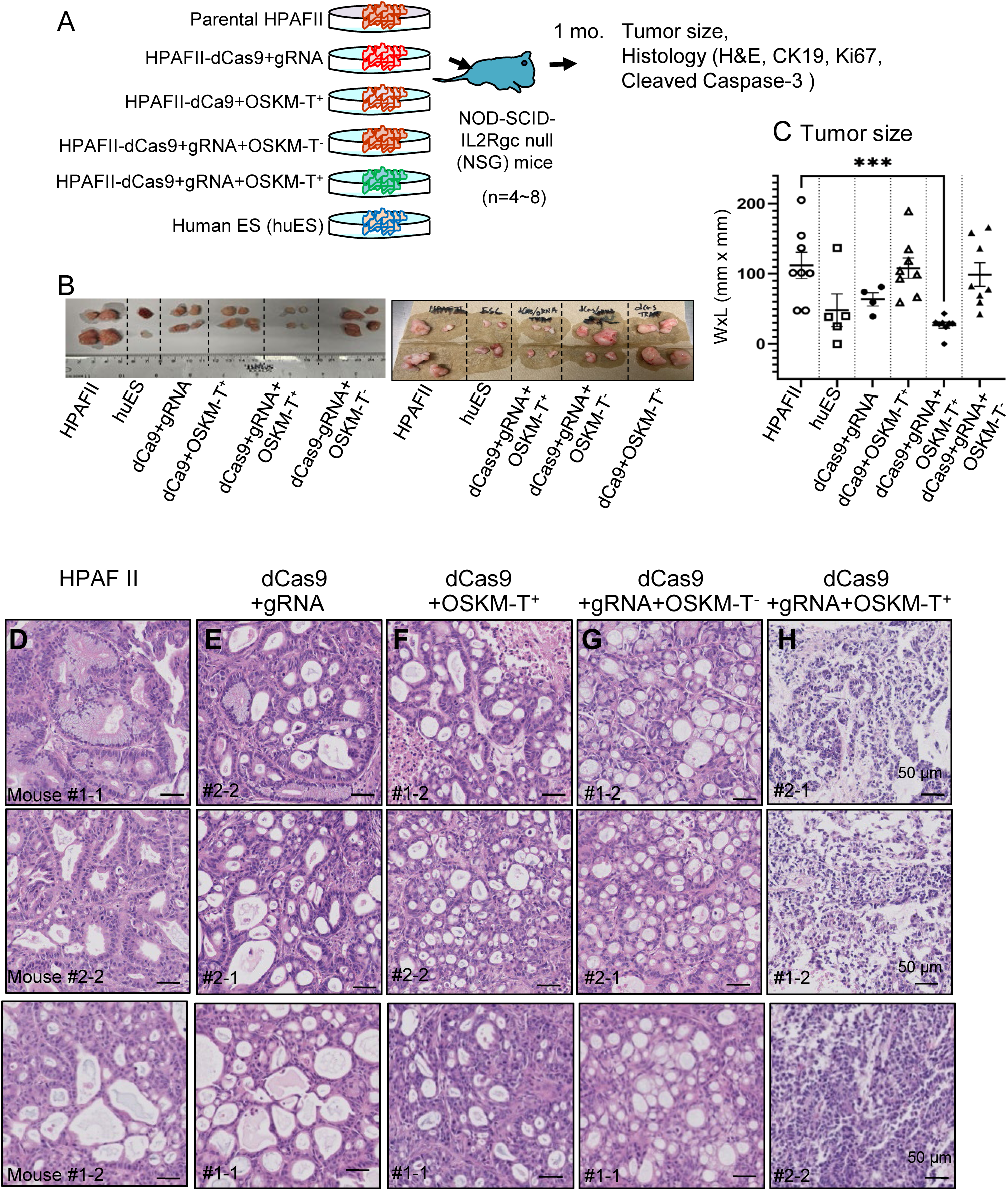
Xenograft Tumor Growth Is Reduced in PDAC Cells Reprogrammed with OSKM and lncRNA Depletion. A. Schematic diagram of xenograft assay of TRA-1-60^+^ and TRA-1-60^−^cells derived from OSKM-reprogrammed HPAFII-dCas9/gRNA, along with controls. Tumors were harvested from NSG mice one month after being subcutaneously injected with dCas9+gRNA+OSKM-T^−^ and dCas9+gRNA+OSKM-T^+^, along with controls: Parental HPAFII, huES, dCas9+gRNA, dCas9+OSKM-T^+^ (n=4-8 for each cell type). B. Representative images of tumors from two independent experiments; two to four mice for each cell type were used in each experiment. C. Comparison of tumor size among different cell types. The differences in tumor sizes (width x height) were tested pairwise using a non-parametric Mann-Whitney test. Only tumor sizes between parental HPAFII and dCas9+gRNA+OSKM-T^+^ are significantly different (*** p=0.0002, n=8 each, Mann-Whitney test). D-H. Representative H&E images of the indicated xenografted tumor samples (D; HPAFII, E; dCas9+gRNA, F; dCas9+OSKM-T^+^, G; dCas9+gRNA+OSKM-T^−^, H; dCas9+gRNA+OSKM-T^+^) from three independent mice. Mouse identifiers (e.g., #1-1, #1-2, #2-1, #2-2) denote individual animals associated with each cell type. The scale bar indicates 50 μm.

**Figure 5.**
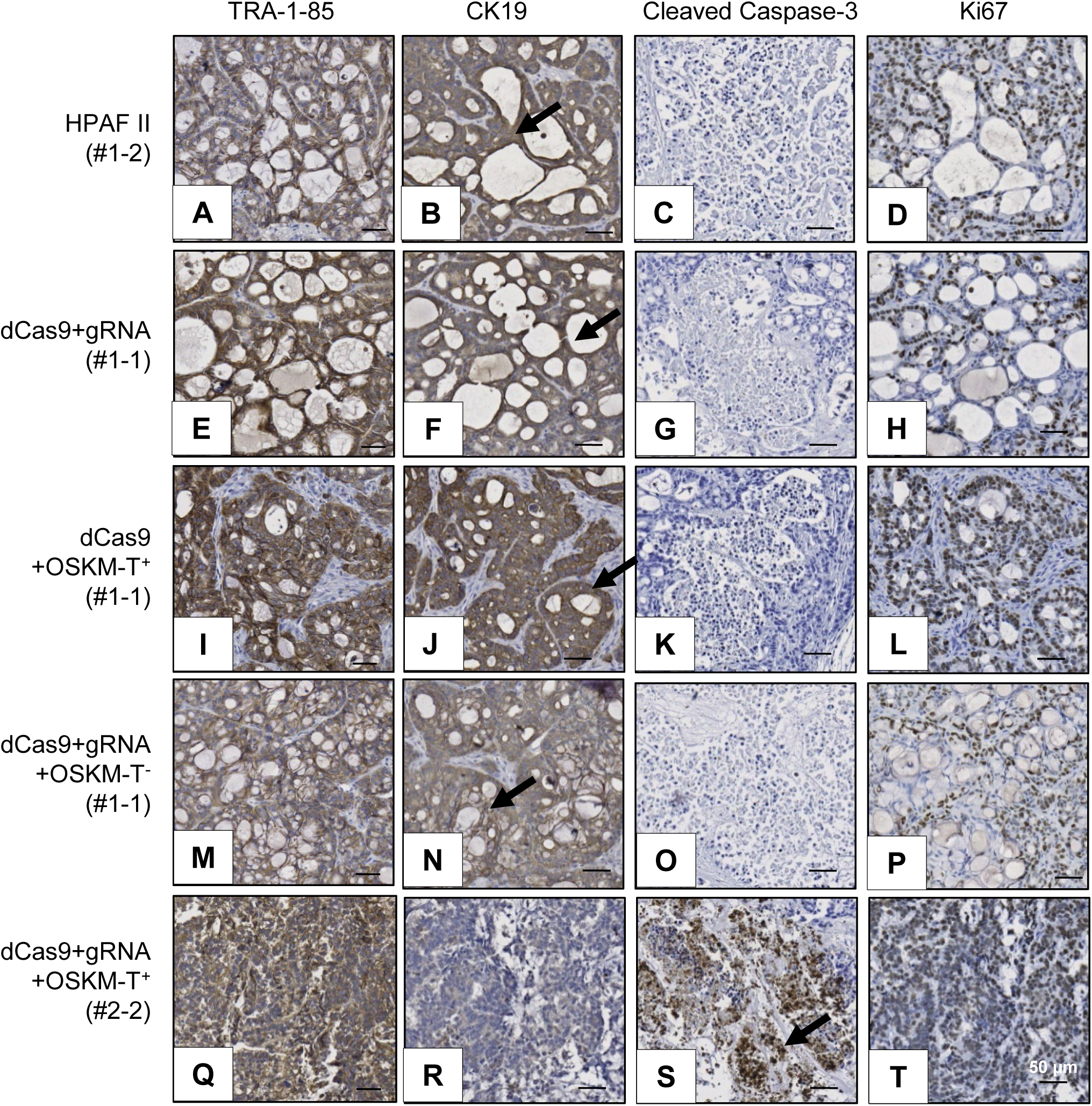
Xenograft Tumors from OSKM/lncRNA-Depleted PDAC Cells Lose Ductal Identity and Express Apoptotic Markers. Representative images of TRA-1-85 human specific marker, human CK19 pancreatic ductal marker, Cleaved Caspase-3 apoptotic marker, and Ki67 proliferation marker staining in xenograft tumors derived from parental HPAFII cells (A–D), dCas9+gRNA cells (E–H), dCas9+OSKM-T+ cells (I–L), dCas9+gRNA+OSKM-T-cells (M–P), and dCas9+gRNA+OSKM-T+ cells (Q–T). Mouse identifiers (e.g., #1-1, #1-2, #2-1, #2-2) denote individual animals associated with each cell type. Arrows indicate positive brown signals. Scale bar: 50 μm.

In histopathology analysis, we confirmed classical grade 2 adenocarcinoma morphology in tumors from all control groups, with strong human-specific marker TRA-1-85, pancreatic ductal marker CK19, and proliferation marker Ki67 immunostainings and absence of apoptotic marker Cleaved Caspase-3 (CC3) immunostaining (Fig. 4D-G, 5A-P). We observed extensive central necrosis, consistent with advanced PDAC pathology. In contrast, tumors derived from dCas9+gRNA+OSKM-T^+^ cells lacked typical adenocarcinoma features, lacking CK19 and showing robust CC3 immunostaining (Fig. 4H, 5Q-T), suggesting a transition toward a less aggressive phenotype.

In all, our analysis underscores that neither OSKM expression nor lncRNA knockdown alone was sufficient to reprogram the malignant phenotype. At the one-month time point, huES-derived tumors were small, composed predominantly of neuronal epithelial cells^90, 91^, which are distinct from HPAFII-derived tumors (Fig. S5A-E). Together, these findings demonstrate that lncRNA depletion in combination with OSKM expression effectively reprogrammed PDAC cells *in vivo*, resulting in reduced tumorigenicity, loss of classical adenocarcinoma identity, and reactivation of apoptotic programs. Importantly, this phenotypic reversal was not achieved by OSKM expression or lncRNA knockdown alone, highlighting the requirement of both to overcome the malignant PDAC state.

### Identification of *ATXN7L3-AS1*, *AC079921.2*, and *INTS4P2* as Cancer-Associated lncRNAs That Restrict OSKM Reprogramming

To identify lncRNAs that maintain PDAC gene expression and are selectively depleted during successful reprogramming, we sequenced genomic DNA from FACS-sorted dCas9–gRNA controls, and gRNA+OSKM-T^+^ and gRNA+OSKM-T^−^ cells (Fig. 6A, Table S5A–E). After normalizing gRNA counts from two independent cultures using Z-score and MAGeCK negative control normalization^92^, we retained lncRNAs with Z-scores > 2 and positive β-scores (*p* < 0.05) in both cultures (Fig. 6A and S5F-H, Table S5F–K, methods).

**Figure 6.**
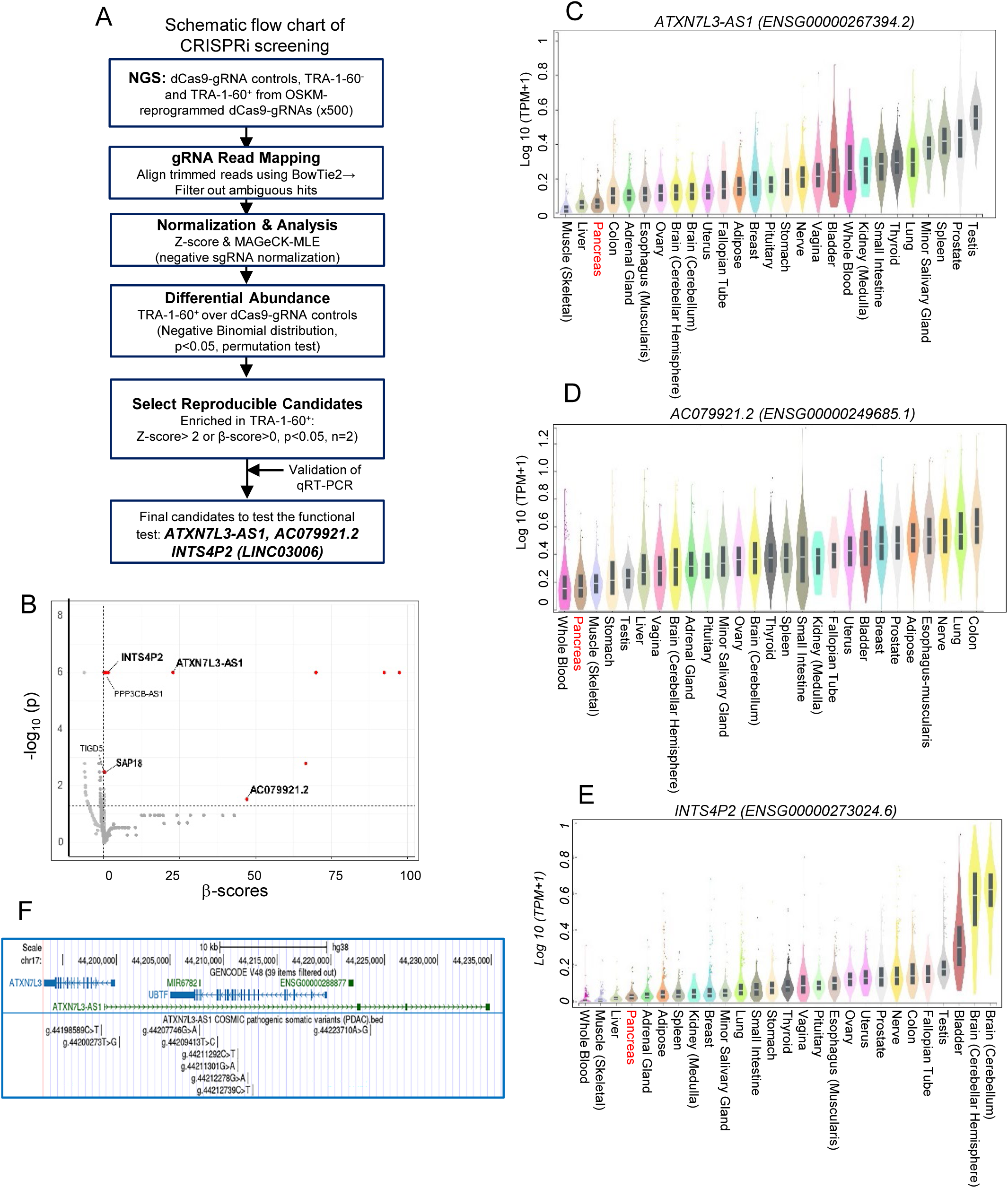
Identification of *ATXN7L3-AS1, AC079921.2, and INTS4P2* as Candidate Barriers to Reprogramming in OSKM/lncRNA-Depleted PDAC Cells. A. Schematic flow chart of CRISPRi screening. B. Significantly depleted lncRNAs in HPAFII-dCas9+gRNA+OSKM-T^+^ cells in comparison to the HPAFII-dCas9-gRNA reference sample are visualized in the volcano plot. The enrichment score of gRNAs is shown as the “β-scores.” Positive and negative β-scores indicate positive and negative selections of gRNAs in the TRA-1-60^+^ compared to the reference condition, respectively. The X-axis shows the β-score of TRA-1-60^+^ compared to the reference. The Y-axis displays –log10 (p-values). The p-value was calculated by ranking lncRNAs with a skewed distribution compared to a uniform null model and permuting to calculate the significance of the skew in distribution. β-score and p-values were calculated separately from two independent experiments and combined to display the top-ranked lncRNAs in one volcano plot. The horizontal and vertical lines indicate the cutoff threshold. The red dots indicate lncRNAs with a β-score over zero and a permutation p-value <0.05. The candidate lncRNAs that were validated by qRT-PCR were highlighted with names. C-E. Expression level of candidate lncRNAs in human normal tissues (GTEx)^91^. Expression values are in TPM (Transcripts per Million), calculated from a gene model with isoforms collapsed to a single gene. The X-axis shows log_10_(TPM+1). Box plots are shown as median and 25th and 75th percentiles; points are displayed as outliers if they are above or below 1.5 times the interquartile range. Pancreatic tissues (n = 362) are highlighted in red. F. UCSC genome browser view of the *ATXN7L3-AS1* locus (chr17:44,198,841–44,234,868; hg38), located between the *ATXN7L3* and *UBTF* genes. Pathogenic somatic variants identified in PDAC from the COSMIC database are displayed along the locus.

This stringent analysis identified several lncRNAs-targeting gRNAs enriched in gRNA+OSKM-T^+^ cells (Fig. 6B). Among these, *ATXN7L3-AS1* (ENSG00000267394), *AC079921.2* (ENSG00000249685), and *INTS4P2* (ENSG00000273024, within LINC03006:ENSG00000290926) were validated by qRT-PCR and found to be significantly downregulated in HPAFII-dCas9+gRNA+OSKM-T^+^ cells compared to control HPAFII-dCas9 cells (Fig. S6A). Importantly, these lncRNAs were minimally expressed or absent in normal pancreatic tissue^93^ (Fig. 6C-E) and remained unchanged after OSKM induction alone (Fig. S6B).

### *ATXN7L3-AS1* Occupies Both PRC2-Poised Developmental and Active Oncogenic Gene Programs in PDAC

To assess functional properties of candidate lncRNAs, we targeted *ATXN7L3-AS1*, *AC079921.2*, and *INTS4P2* with CRISPR-dCas9 in PDAC cells (HPAFII and CAPAN1) and normal ductal cells (H6c7). Knockdown of *ATXN7L3-AS1*, *AC079921.2*, and *INTS4P2* reduced expression of PDAC-enriched genes, including basal markers (*KRT5, TP63*) and the lineage factor GATA6, while modestly increasing progenitor markers (*PDX1, FOXA2, SOX9*) upon depletion of *ATXN7L3-AS1* and *AC079921.2* (Fig. S6C). Importantly, depletion of *ATXN7L3-AS1* and *AC079921.2* had the opposite effect on normal ductal H6c7 cells, supporting a cancer-specific role for these lncRNAs (Fig. S6D). Notably, the *ATXN7L3-AS1* locus harbors recurrent pathogenic mutations across multiple cancers, underscoring its potential role in establishing and maintaining malignant states (OSMIC^94^; Fig. 6F, Table S6A-B).

Chromatin Isolation of RNA purification (ChIRP-seq)^95^ analysis identified that *ATXN7L3-AS1* primarily localizes to CpG island promoters, enriched for ZNF transcription factor motifs, of genes involved in RNA Pol II regulation, actin cytoskeletal remodeling, and cell migration, including neuronal migration (Fig. 7A–D, S7A–B, Table S6C–D). Notably, *ATXN7L3-AS1*– occupied genes overlapped with binding sites of UBTF, ZBTB7A, and PRC2 components (SUZ12, EZH2), as well as regions marked by H3K27me3 (Fig. 7D, S7C–E).

**Figure 7.**
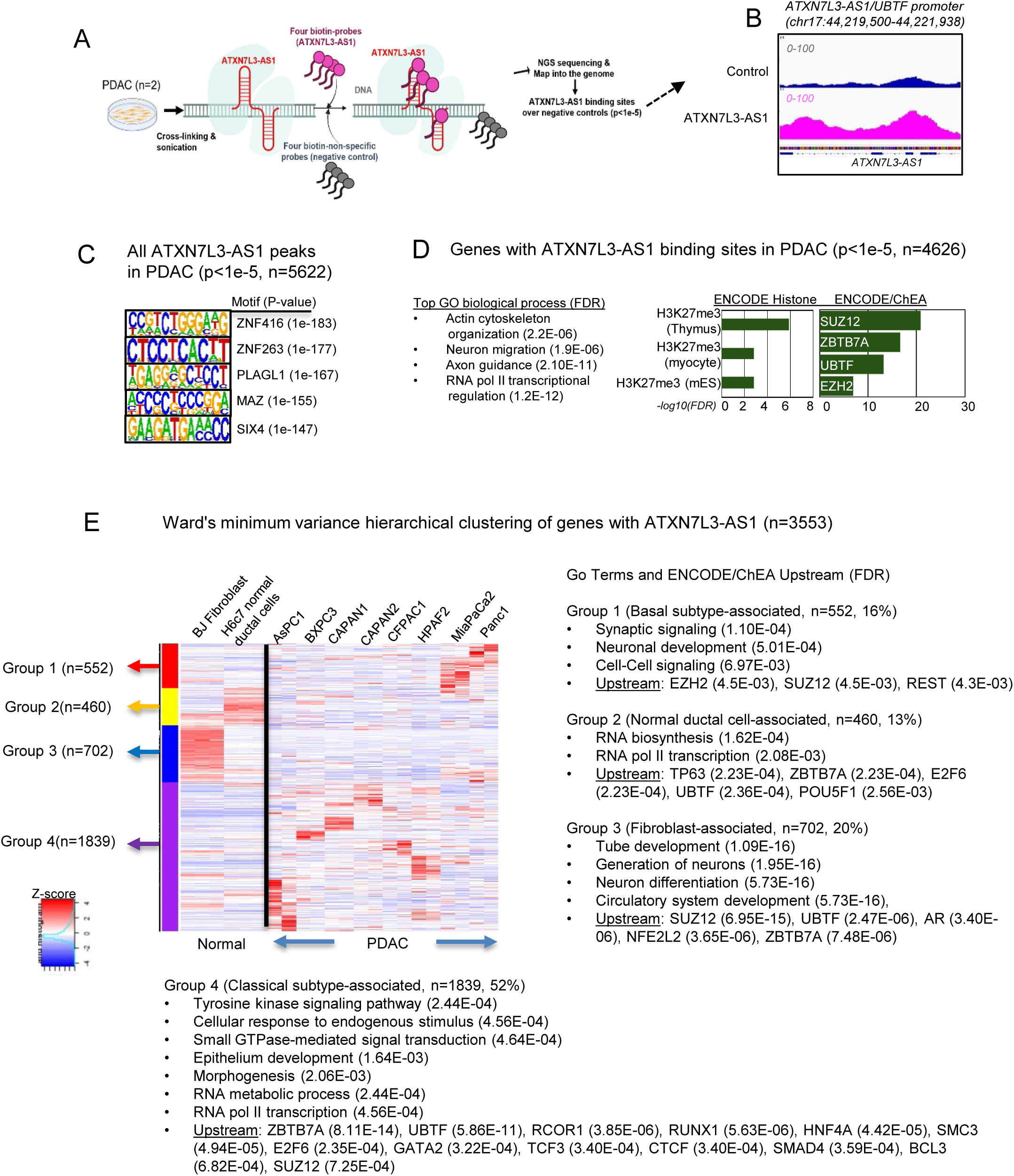
*ATXN7L3-AS1* Binds Two Categories of Genes in PDAC: Developmental Programs Such as Cell Migration, Which It Preserves in a Poised State, and Cancer-Associated Genes, Which It Maintains in an Active State. A-C. Mapping *ATXN7L3-AS1* binding sites using ChIRP-seq. (A) Schematic of ChIRP-seq strategy for mapping *ATXN7L3-AS1* genomic binding sites in PDAC cells (HPAFII and CAPAN1, n=2 each cell). *ATXN7L3-AS1*–RNA/DNA complexes were cross-linked, captured with biotinylated antisense probes, and sequenced; non-specific probes served as controls. (B) Representative genome browser tracks of *ATXN7L3-AS1* ChIRP-seq peaks. *ATXN7L3-AS1* binding is enriched at the *UBTF* promoter, which lies within the *ATXN7L3-AS1* genomic locus. (C) HOMER motif analysis of the sequence of *ATXN7L3-AS1* peaks. De novo motif discovery was performed with HOMER on all 5,622 *ATXN7L3-AS1* ChIRP-seq peaks (p<1e-5). Significantly enriched motifs are shown, ranked by –log₁₀(p). D-E. Genes with *ATXN7L3-AS1* Peaks. (D) Functional enrichment of genes with *ATXN7L3-AS1* binding peaks (p < 1e–5), including representative GO terms, ENCODE histone mark annotations, and ENCODE/ChEA-predicted regulators. (E) Gene signatures associated with *ATXN7L3-AS1*–bound genes. Ward minimum variance hierarchical clustering of genes with *ATXN7L3-AS1* across normal and PDAC cell lines. The heatmap indicates Z-score expression. Representative Go categories and ENCODE/ChEA Consensus upstream analysis of genes in each group (FDR<0.05).

Next, we examined the expression of *ATXN7L3-AS1*–occupied genes across normal and PDAC cells^96^ (Fig. 7E, Table S6E-H). Ward’s clustering analysis^97^ partitioned these genes into four transcriptional modules with distinct biological signatures: a fibroblast-specific cluster (Group 3, 20%) enriched in developmental programs such as neuronal differentiation and tube formation; a normal ductal cell–associated cluster (Group 2, 13%) dominated by RNA metabolism and transcriptional regulation; a basal subtype–associated cluster (Group 1, 16%) enriched in synaptic signaling and neuronal differentiation, consistent with neuronal mimicry of basal subtypes; and a broad classical subtype–associated cluster (Group 4, 52%) enriched in RNA metabolism, tyrosine kinase signaling, small GTPase pathways, and epithelial morphogenesis (Fig. 7E, Table S6E-H). Upstream regulator analysis revealed cluster-specific differences: SUZ12/PRC2 targets were preferentially enriched in the fibroblast- and basal subtype–associated groups (Groups 3 and 1), consistent with Polycomb-maintained developmental programs. By contrast, UBTF emerged as the top regulator in the broad classical cluster (Group 4), where SUZ12 ranked lower (Fig. 7E).

Consistent with this model, *ATXN7L3-AS1*–bound tubular ductal epithelial development genes are normally silenced by PRC2/H3K27me3 but remain active in PDAC (Fig. 8A–B). Similarly, OSKM-aberrantly upregulated genes, which significantly overlap with *ATXN7L3-AS1*– bound genes, were particularly enriched in migration and morphogenesis pathways (Fig. 8C). Together, these findings suggest that *ATXN7L3-AS1* lncRNA sustains malignant PDAC identity by maintaining epithelial cancer–associated programs in an active state while keeping PRC2-target developmental genes poised, thereby supporting both plasticity and malignant transcriptional memory.

**Figure 8.**
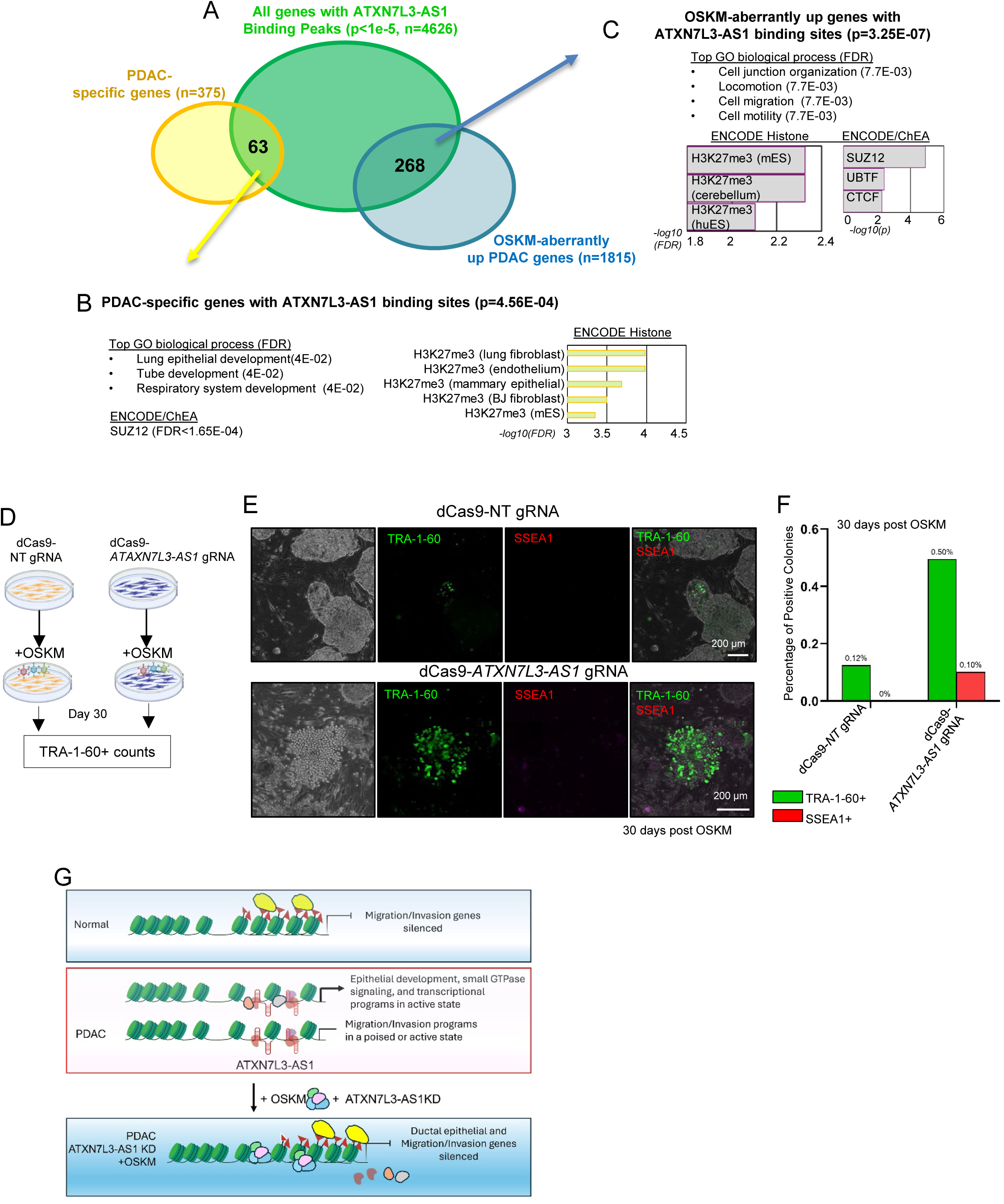
*ATXN7L3-AS1* Binds Malignant Genes and Impedes OSKM Reprogramming in PDAC. A. Intersection of genes with *ATXN7L3-AS1* binding sites with PDAC-specific genes and OSKM-aberrantly upregulated genes (p=4.56E-04, 3.25E-07, respectively, hypergeometric test). B-C. Enriched GO categories, histone modifications, and upstream regulators associated with *ATXN7L3-AS1* peaks overlapping (B) PDAC-specific genes or (C) OSKM-aberrantly upregulated genes. D-F. Reprogramming of PDAC cells depleting *ATXN7L3-AS1* lncRNAs. (D) Schematic of CRISPRi-based depletion of *ATXN7L3-AS1* during OSKM induction in PDAC cells. (E) Representative images of TRA-1-60⁺ colonies derived from HPAFII-dCas9–*ATXN7L3-AS1* gRNA cells at day 30 of OSKM reprogramming; HPAFII-dCas9–NT gRNA cells serve as controls. (F) Quantification of colony phenotypes showing the percentage of TRA-1-60⁺ (green) or SSEA1⁺ (red) colonies among total colonies counted. G. Model of *ATXN7L3-AS1* function in sustaining malignant PDAC transcriptional programs. In normal ductal epithelial cells, migration and invasion programs are silenced. In PDAC, *ATXN7L3-AS1* associates with CpG island promoters to maintain epithelial development, small GTPase signaling, and transcriptional programs in an active state, while keeping migration and invasion pathways in a poised state. Upon *ATXN7L3-AS1* knockdown during OSKM reprogramming, this protection of the cancer genome is lost, allowing OSKM to silence PDAC identity gene programs and thereby enable reprogramming.

Similarly, *AC079921.2* was associated with the expression of invasive hallmark PDAC genes involved in extracellular matrix (ECM) remodeling (Fig. S8A–C, Table S7) across TCGA Pan-Cancers. OSKM-unresponsive or aberrantly upregulated genes overlapping with these sets were likewise enriched in morphogenesis, migration, and predicted SUZ12 targets (Fig. S8D–E). Thus, *AC079921.2* reinforces the malignant identity of PDAC by sustaining pro-invasive and morphogenesis programs resistant to OSKM.

Finally, functional experiments further validated that *ATXN7L3-AS1* depletion markedly enhanced OSKM-mediated reprogramming in PDAC cells, but not in normal ductal cells (Fig. 8D–F), underscoring its cancer-specific role in maintaining malignant transcriptional memory. Collectively, these findings reveal that *ATXN7L3-AS1* and *AC079921.2* sustain malignant PDAC programs by engaging both PRC2-regulated developmental pathways and active promoters of cancer-associated genes, thereby reinforcing plasticity and malignant memory (Fig. 8G).

## Discussion

OSKM-mediated reprogramming can reset cell identity in normal and aged cells by remodeling the epigenome, yet most solid tumors remain largely refractory^13–19^. This resistance has long remained an enigma, given the well-known plasticity of cancer cells and their expression of pluripotency factors. Here, we optimized a high-efficiency, non-integrating gene delivery system^60^, amplified primary PDAC cells^59^, and developed an analytical framework to account for inter-sample heterogeneity. While normal pancreatic ductal epithelial cells are intrinsically more resistant to reprogramming than fibroblasts or acinar cells, they can ultimately be reprogrammed with sustained expression of OSKM. In contrast, under the optimized conditions, we found that PDAC cells failed to undergo full reprogramming, in part due to the aberrant activation of PRC2-target developmental genes, including those involved in ECM remodeling and mesodermal lineage programs. A subsequent CRISPRi screen revealed that lncRNAs act as major gatekeepers of malignant identity, driving resistance to reprogramming. Because lncRNAs often act as scaffolds for PRC2 and are expressed at low levels, conventional RNA-seq pipelines may fail to detect those functionally involved in preserving malignant memory. Whether oncogenic KRAS directly contributes to reinforcing such lncRNA-mediated epigenetic barriers is an important question for future studies.

### *ATXN7L3-AS1* and *AC079921.2* as Gatekeepers of PDAC Identity

LncRNAs can contribute to transcriptional memory by shaping chromatin architecture and regulating gene expression through chromatin-modifying enzymes^38–49^. Our data suggest that *ATXN7L3-AS1* acts as a dual regulator of transcriptional memory in PDAC. By binding CpG-rich promoters enriched for ZNF TF binding motifs, it preserves PRC2-target developmental programs such as ECM remodeling and cell migration in a poised, while simultaneously maintaining activation of epithelial cancer-associated genes. This dual activity enables PDAC cells to sustain both latent plasticity and active malignant transcriptional programs, providing a mechanistic basis for their resistance to reprogramming. This is consistent with prior work reporting that lncRNAs scaffold chromatin modifiers preserve lineage fidelity^91–94^. Importantly, *ATXN7L3-AS1* loci harbor pathogenic PDAC mutations (Fig. 6F) and *UBTF*, a known oncogenic TF in other cancers^96–98^, suggesting that its role in PDAC spans both structural and regulatory mechanisms.

By contrast, *AC079921.2* expression was correlated with ECM remodeling, cell adhesion, and migration genes—hallmarks of mesenchymal-like and invasive phenotypes (Fig. S8A-C). Like *ATXN7L3-AS1*, these gene sets were enriched for PRC2 targets, suggesting that *AC079921.2* antagonizes PRC2-mediated repression. Supporting this view, genes aberrantly upregulated or unresponsive to OSKM reprogramming significantly overlapped with *AC079921.2*–correlated genes in PDAC (Fig. S8D-I). Thus, *AC079921.2* sustains a mesenchymal-like, invasion-prone transcriptional program that resists reprogramming while promoting adaptability to microenvironmental cues. Whether *ATXN7L3-AS1* or *AC079921.2* physically interacts with PRC2 to modulate its activity is unknown. Together, these findings establish *ATXN7L3-AS1* and *AC079921.2* as gatekeepers of PDAC identity, acting through distinct yet complementary mechanisms to preserve transcriptional memory and plasticity, thereby stabilizing malignant states.

### Combining lncRNA Depletion with OSKM Induction is Required to Erase PDAC Identity

While knockdown of *ATXN7L3-AS1*, *AC079921.2*, or *INTS4P2* moderately reduced PDAC marker expression, neither lncRNA depletion nor OSKM induction alone was sufficient to fully erase PDAC identity or suppress malignancy *in vitro* or *in vivo* (Figs. 3-5). In xenografts, lncRNA depletion modestly reduced tumor size, likely due to decreased proliferation without increased apoptosis, but did not induce reprogramming or erase adenocarcinoma features. Similarly, OSKM expression yielded tumors with more differentiated histology and reduced proliferation, yet these tumors retained core transcriptional programs characteristic of PDAC.

In contrast, combining lncRNA knockdown with OSKM induction (dCas9+gRNA+OSKM-T^+^ cells) erased adenocarcinoma identity, reactivated apoptotic pathways, and markedly impaired tumorigenesis *in vivo*. Under these conditions, OSKM converted cells into NANOG⁺ pluripotent-like states in which TSGs were robustly reactivated. Notably, this requirement for pluripotent conversion to enable TSG re-expression is distinct from other reprogramming contexts, where NANOG⁺ states are often linked to malignant potential. Together, cancer-associated lncRNAs emerge as key determinants of malignant persistence, whose suppression is necessary to unlock the reprogramming potential of OSKM and redirect it away from malignancy.

### Toward a new reprogramming cocktail: lncRNA depletion sensitizes PDAC to OSKM-induced cell fate transitions

Our prior work demonstrated that full reprogramming of PDAC cells into a pluripotent state was impeded, with partially reprogrammed states requiring sustained ectopic OSKM expression^16^. Herein, we extend these findings by proposing that lncRNA depletion sensitizes PDAC cells to OSKM-mediated reprogramming by weakening epigenetic barriers that preserve malignant identity. To enhance clinical relevance, we optimized a non-integrating, Sendai virus–mediated system for transient OSKM expression across multiple patient-derived PDAC cells^60^.

Beyond pluripotency, lineage reprogramming of cancer cells into alternative fates—such as dendritic cells—has been demonstrated^98–100^. Whether lncRNA depletion similarly sensitizes PDAC cells to acquire non-permissive cell states^101–103^ remains an open question. Targeting lncRNAs enforcing transcriptional memory could enhance responsiveness to reprogramming cues and improve lineage fidelity.

### Conclusions

Here, we integrate prior work on OSKM reprogramming with PRC2 biology and lncRNA-mediated transcriptional memory to explain how PDAC cells can simultaneously be plastic and rigid. Specifically, we show that cancer-associated lncRNAs act as gatekeepers of malignant identity, preserving ductal epithelial and invasion-prone programs that resist reprogramming. Targeting these lncRNAs removes this barrier, unlocking the capacity of OSKM to erase malignant identity, reactivate tumor suppressor pathways, and impair tumorigenesis. This conceptual framework establishes malignant memory as an actively maintained epigenetic state and suggests that lncRNA-directed therapies may sensitize tumors to differentiation- and reprogramming-based therapeutic interventions in pancreatic cancer.

### Limitations of the Study

Our study focuses on PDAC-specific lncRNAs that sustain the cancer identity and utilizes OSKM to induce the reprogramming of PDAC toward pluripotency. However, lncRNAs that sustain the cancer identity may differ across cancer types, and the effective reprogramming into other lineages will likely require tailored combinations of lineage-specific transcription factors.

## RESOURCE AVAILABILITY

### Lead Contact

Further information and requests for resources and reagents should be directed to and will be fulfilled by the lead contact, Jungsun Kim.

### Materials availability

- Plasmids, PDX tumors, and reprogrammed cancers generated in this study will be available upon request.
- This study did not generate new unique reagents.

### Data and code availability

- Raw data and codes for sequencing data have been deposited at NCBI GEO and are publicly available as of publication. Accession numbers will be listed on the key resource table. The lead contact, upon request, will share any raw data reported in this paper.
- Codes for sequencing data have been deposited on GitHub and are publicly available as of publication.
- Any additional information required to reanalyze the data reported in this paper is available from the lead contact upon request.

## ACKNOWLEDGMENTS

We thank Drs. Ken Zaret and Theresa Lusardi for comments on the manuscript. We thank Dr. Jeffrey Drebin for providing de-identified resected tumor specimens for the initial phase of this study, which was exempt from IRB review. We thank the Brenden-Colson Center for Pancreatic Care, the Oregon Pancreas Tissue Registry, the OHSU Knight Biostatistics Shared Resource and Biolibrary (NCI P30CA69533), and the Molecular Virology Core at the Oregon National Primate Research Center (NIH P51OD011092). The IRB protocol is the Oregon Pancreas Tissue Registry (IRB #3609). The work was funded by the CRUK-OHSU Project Award (2018-CRUK-OHSU-001 to JK), MRF New Investigator Grant (GCNCR1042A to JK), and Knight CEDAR (68182-906-000, 68182-933-000, 68182-939-000, 68182-954-000, Manuscript Prep 2024-1978) from the Cancer Early Detection Advanced Research Center at OHSU Knight Cancer Institute. Life Science Editors edited this article.

## AUTHOR CONTRIBUTIONS

Conceptualization, JK; Methodology, DG, SL, CMF, SR, MB, AH, TE, EM, JT, MCC, JL, and WY; Validation, DG, SL, CMF, SR, SF, TM, and WY; Formal Analysis, DG, SL, CMF, SR, AH, SF, TM, WY, EF, ST, and JK; scRNAseq analysis supervision, ST and JK; Investigation, D.G., SL, CMF, SR, MB, SF, TM, WY, EF, ST, and JK.; Resources, JL, DK, BCS, EF, and RCS; Data Curation, DG, SL, CMF, SR, AH, SF, and ST; Writing-original draft, JK; Writing-Review & Editing, DG, SL, CMF, SR, and JK; Visualization, DG, CMF, SR, AH, SF, and ST, Supervision, JK; Project Administration, JK; Funding Acquisition, JK.

## DECLARATION OF INTERESTS

The authors declare no competing interests.

## METHODS

### Derivation of Patient-Derived Xenografts (PDX) and Tissue Culture

De-identified human PDAC specimens were obtained under the University of Pennsylvania study (IRB-exempted) or the Oregon Pancreas Tissue Registry protocol (IRB00003609). Written informed consent was obtained from all participants. The OHSU Institutional Review Board approved all experimental protocols. All methods were carried out in accordance with relevant guidelines and regulations. All animal work for PDX tumors was performed with the OHSU Institutional Animal Use and Care Committee (IACUC) approval. The derivation, expansion, culture, and characterization of PDAC PDXs, are described in our protocol paper^59^. Culture conditions for H6c7 and other PDAC cell lines are described^59^. BJ6 fibroblasts were cultured in DMEM supplemented with 10% fetal bovine serum (FBS)^3^.

### Reprogramming of Pancreatic Cancer Cells and Normal Controls

Reprogramming was performed using the CytoTune-iPS Sendai Reprogramming Kit (Invitrogen), which delivers the non-integrating Sendai virus encoding OCT4, SOX2, KLF4, and c-MYC (OSKM)^104^. Viral titers for Sendai virus and lentivirus were described in our earlier publication^60^. On day 7 post-transduction, cells were harvested for RNA sequencing and re-seeded onto irradiated mouse embryonic fibroblasts (MEFs) to support further reprogramming until 30 days.

### CRISPR interference (CRISPRi) Screen

We performed CRISPRi–based screens targeting 691 cancer-associated lncRNAs using a pooled lentiviral library containing 7,087 gRNAs (10 gRNAs per lncRNA, plus non-targeting controls, Millipore, Table S3A)^83, 84^. The HPAFII and H6c7 cells expressing dCas9-KRAB were generated by transducing HPAFII and H6c7 cells with lentiviruses carrying the dCas9-Bsr gene at an MOI of 1, followed by selecting cells for Blastidin. The CRISPRi efficiency was assessed in the clonal HPAFII and H6c7 cells expressing dCas9 with positive controls. The clonal HPAFII-dCas9 and H6c7-dCas9 were subsequently transduced with a lentiviral library of 7087 sgRNAs at an MOI of 0.1-0.3 with 300-500x coverage to maintain an average 300-500-fold representation of each sgRNA. After puromycin selection, cells were collected for reference controls (300x, 500x). HPAFII and H6c7 cells were reprogrammed with OSKM Sendai viruses using the OSKM Sendai viruses^104^. The CRISPRi-targeted cells were always maintained at 300-500 coverage throughout the procedure.

For downstream analysis, cells were harvested after 30 min incubation with AccuMAX and processed using a mouse cell depletion kit as previously described^59^. For flow cytometry analysis or sorting, DNase I (0.1 mg/ml) was added to prevent cell aggregation, and TRA-1-85⁺ expression was used to identify human cells, while BFP marked gRNA-expressing cells.

### RNA-seq Data Processing and Normalization

Bulk RNA-seq was performed as previously described^60, 105^. Raw FASTQ files were processed using fastp to generate clean reads by removing adapter sequences, reads containing poly-N, and low-quality reads. During this step, quality metrics such as Q20, Q30, and GC content were also calculated. All downstream analyses were conducted using the high-quality clean reads. The reference genome and gene annotation files were downloaded directly from the genome database. The genome index was built using HISAT2 (v2.0.5), and paired-end clean reads were aligned to the reference genome using HISAT2 (v2.0.597)^106^. FeatureCounts (v1.5.0-p3) were used to quantify the number of reads mapped to each gene. FPKM (Fragments Per Kilobase of transcript per Million mapped reads) values were then calculated for each gene based on its length and the number of mapped reads as needed. Differential expression analysis was performed using the DESeq2 R package (v1.20.0), which employs statistical routines based on the negative binomial distribution to identify DEGs from count data^76^. P-values were adjusted for multiple testing using the Benjamini-Hochberg method to control FDR. Genes with an adjusted P-value ≤ 0.05 were considered significantly differentially expressed. RNA-seq data are available on NCBI GSE247243.

### Identification of PDAC Gene Signatures Repressed by OSKM

To identify PDAC-specific gene categories, we first performed differential expression analysis using DESeq2 to compare each PDAC cell to normal controls (BJ fibroblasts and H6c7 pancreatic epithelial cells). Genes that were significantly upregulated in PDAC cells (FDR < 0.05, log₂ FC > 1) were selected. We then intersected the upregulated gene sets across all four PDAC samples to define a core set of PDAC-upregulated genes (Table S2A). To assess the impact of OSKM induction, we compared gene expression profiles before and after OSKM treatment and identified differentially expressed genes using the same thresholds (FDR < 0.05, log₂FC> 1 or < –1, Table S2B).

### Pattern Classification Following OSKM and Gene Ranking by Weighted Hamming Distance

DEGs before and after OSKM induction (day 7 vs. day 0) were identified for each cell type using the LRT implemented in DESeq2^76^. To establish robust thresholds for DEG calling, we evaluated how closely observed DEG statistics matched the expected null hypothesis distribution. Specifically, we constructed empirical quantile-quantile (Q-Q) plots comparing DEG statistics from genes presumed to be non-differentially expressed (those with simultaneously high adjusted p-values and low log2(FC)) against their expected null distribution. We applied Huber regression to these Q-Q plots and measured the mean absolute deviation (MAD) between the fitted regression line and the identity line over the range of observed statistics. Higher MAD values indicate greater divergence from the null distribution, reflecting genuine biological signals. MAD values were computed across a range of adjusted p-values and log2(FC) threshold combinations and visualized as a heatmap. Based on this analysis, genes with log2(FC) > 0.2 and FDR < 0.05 were considered significantly differentially expressed, as these thresholds corresponded to a clear elevation in MAD.

To classify gene expression responses to OSKM induction, we analyzed DEGs across six cell types—two normal controls and four PDAC cells ([normal#1][normal#2][cancer#1][cancer#2][cancer#3][cancer#4])—using six predefined expression patterns: i) Downregulated in normal, unchanged in cancer: [−1][−1][0][0][0][0], ii) Downregulated in normal, upregulated in cancer: [−1][−1][1][1][1][1], iii) Downregulated in normal, upregulated or unchanged in cancer: [−1][−1][0/+1][0/+1][0/+1][0/+1], iv) Unchanged in normal, upregulated in cancer: [0][0][1][1][1][1], v) Upregulated in normal, downregulated in cancer: [1][1][−1][−1][−1][−1], vi) Upregulated in normal, unchanged in cancer: [1][1][0][0][0][0]. These patterns were used to stratify gene responses to OSKM and identify cancer-specific regulatory behaviors. We classified genes in Class I ([−1][−1][0][0][0][0]) as OSKM-unresponsive genes, and genes in Class II ([−1][−1][1][1][1][1]), III ([−1][−1][0/+1][0/+1][0/+1][0/+1]), and IV [0][0][1][1][1][1] as OSKM-aberrantly upregulated genes (Table S2F-H).

To rank genes within each predefined expression pattern category, we used a weighted Hamming distance metric^77^ relative to the query vector. Let q denote the query vector representing a specific regulation profile (e.g., upregulated in normal with minimal change in cancer samples), and let cat(S) denote the categorical expression vector of a gene. The weighted Hamming distance d is calculated as: d = ∑(i=1 to N) w_i_ × Ind(q_i_ ≠ cat(S)_i_) where N is the number of conditions or replicates, w_i_ is the weight assigned to the *i*th condition, and Ind is the indicator function that returns 1 if the query and gene vector differ at position *i*, and 0 otherwise. This metric penalizes deviations from the target pattern, with weights chosen to balance representation across biological conditions and to reflect the hierarchical structure of replicates within each condition. Genes with lower values of the weighted Hamming distances are considered better matches to the specified query. For each query of interest, genes were ranked in ascending order based on their weighted Hamming distance (Table S2F-H).

### Single-cell RNAseq analysis

Single-cell RNA sequencing (scRNA-seq) was performed using the Chromium Next GEM Single Cell 5’ Kit (v2.0, 10x Genomics). All scRNA-seq samples were processed using Seurat v5. Following quality control, cells were normalized, scaled, batch-corrected, and filtered in accordance with best practices outlined in the Seurat v5 vignette^107^. Cells expressing gRNA were filtered for BFP. After filtering out low-quality cells with >10% mitochondrial gene content and regressing out cell cycle–associated transcripts, we retained high-quality transcriptomes for downstream analysis (Fig. S4A-B). Clustering was performed using the Seurat integration workflow with the CCA (Canonical Correlation Analysis) method, at a resolution of 0.5 using the Louvain algorithm. Harmony integration was also performed to assess potential differences compared to the Seurat CCA-based integration. scRNAseq data set was extracted from human PDAC (n=6, GSE212966^85^), human and mouse normal pancreas (n=4, GSE84133^86^), human early embryo before the three germ layers formed (D6-D14, n=555, GSE136447^108^), and iPSC derived from Parkinson’s patients’ fibroblasts (n=12, GSE183248^109^). “PDAC-specific signature genes” were curated from published scRNA-seq datasets by identifying transcripts enriched in PDAC cells but expressed in fewer than 20% of normal pancreatic cells (Table S4D)^85, 86^. Tumor suppressor genes (TSG) were curated from the OncoKB database^88^ (Table S4E). NT clusters were manually defined based on NANOG and TRA-1-60 expression profiles. Cells were grouped according to their expression of these markers, with clusters curated based on predefined thresholds rather than unsupervised clustering algorithms.

### Next-Generation Sequencing for Identification of Depleted gRNAs in Reprogrammed Cells (Table S5)

Genomic DNA from each group was subjected to nested PCR amplification to generate NGS libraries, enabling identification of enriched or depleted sgRNAs in TRA-1-60⁺ reprogrammed cells compared to either reference controls or TRA-1-60⁻ cells. These analyses were performed on two independently passaged biological cultures, separated by 3–6 passages. The hit count corresponding to each sgRNA was computed from raw FastQ files using Millipore’s custom pipelines. The raw hit counts were normalized by Z-score for exploratory analysis. The MAGeCK package with negative control sgRNA normalization was used to analyze the enriched lncRNAs^92^. The MAGeCK algorithm first performs normalization and then uses information borrowing across genes to estimate variance^92^. A negative binomial model is then fit and used to test for differential hits on lncRNAs across samples. Significance testing used permutations. We used the MAGeCK mle module since there were more than two conditions. The MAGeCKFlute R package was used with R version 4.4 for data analysis and visualization^110^. To ensure robustness, raw gRNA counts from both experiments were merged, scaling the second dataset to correct for batch differences (Exp1/Exp2 ≈ 1.86), with counts rounded up to the nearest integer (Table S5J). gRNAs consistently enriched from two independent samples, as defined by either Z-scores > 2 and positive beta scores (p < 0.05), were retained (Table S5K).

### Chromatin Isolation by RNA Purification followed by sequencing (ChIRP-seq)

CAPANI cells and HPAFII were transfected with an *ATXN7L3-AS1* overexpression plasmid to generate cells with elevated nuclear expression of *ATXN7L3-AS1*. Cells (2 × 10^7^) were washed three times with ice-cold PBS and crosslinked by adding 37% formaldehyde solution (1:36 dilution) for 10 min at room temperature (25 °C). Crosslinking was quenched with 2.5 M glycine (1:20 dilution) for 5 min. Crosslinked cells were scraped, collected by centrifugation at 1,000 × g for 3 min, washed twice with pre-cooled PBS, and pelleted by centrifugation. Cell pellets were resuspended in 1 mL cold lysis buffer per 2 × 10^7^ cells and sonicated (30 s on/30 s off, five cycles) to obtain chromatin fragments. Lysates were clarified by centrifugation at 12,000 rpm for 10 min at 4 °C, and supernatants were stored at −80 °C until ChIRP.

Cell lysates were mixed with RNA–protein hybridization buffer at a 1:2 ratio and incubated overnight at room temperature with 5 μL of biotin-labeled probes tiling the target RNA. Streptavidin magnetic beads (50 μL) were washed three times with hybridization buffer and then incubated with the sample–probe mixture for 4 h at room temperature. Beads were washed five times with 1 mL wash buffer and bound complexes were eluted with 50 μL elution buffer at 37 °C, 1,000 rpm for 5 min. Eluates were treated with proteinase K (2.5 μL) overnight at 65 °C, and DNA was purified using a DNA purification kit. Purified DNA was dissolved in 50 μL DEPC-treated water for downstream library preparation. ChIRP-enriched DNA was used to construct sequencing libraries with the VAHTS mRNA-seq V3 Library Prep Kit. Libraries were prepared by end repair, adaptor ligation, magnetic bead purification, and PCR amplification, followed by stringent quality control. The sequencing was performed on the Illumina platform.

### ChIRP-seq data processing and analysis

Raw sequencing reads were quality-filtered to remove low-quality reads, adaptor sequences, and PCR duplicates. Clean reads were aligned to the human genome (hg38) using STAR2^111^, allowing up to two mismatches. Only uniquely mapped reads were retained for downstream analyses. Peak calling was performed with MACS^112^ using a p-value cutoff of 1e-5, and binding peaks were defined by clustering overlapping reads. Peak characteristics, including genomic distribution, length, spacing, and density, were analyzed. Annotation of *ATXN7L3-AS1* binding sites was performed with HOMER (annotatePeaks.pl) to assign peaks to genomic features (TSS, −1 kb to +100 bp; TTS, −100 bp to +1 kb; CDS exons; 5′ UTR exons; 3′ UTR exons; CpG islands; repeats; introns; and intergenic regions)^113^. Genes associated with significant peaks were subjected to functional enrichment analyses to identify biological pathways potentially regulated by *ATXN7L3-AS1*. Motif enrichment analysis was performed using HOMER.

### Gene Ontology (GO) and Upstream Analysis

GO analysis was performed using Enrichment Analysis^114^ or the ToppGene Suite^115^. To determine whether gene sets of interest are enriched for putative TF targets, we performed ChIP-X enrichment analysis using datasets from ENCODE^116^ or ChEA^117^, accessed through Enrichr^118^ or X2K^119^. ChIP-X refers to genome-wide TF binding data obtained from ChIP-ChIP, ChIP-Seq, ChIP-PET, or DamID experiments^117^. The p-value is computed using Fisher’s exact test or the hypergeometric test. This is a binomial proportion test that assumes binomial distribution and independence for the probability of any gene belonging to any set. The q-value is an adjusted p-value using the Benjamini-Hochberg method for correction for multiple hypothesis testing. The combined score is calculated by taking the logarithm of the p-value from Fisher’s exact test and multiplying it by the Z-score, which reflects the deviation from the expected rank. The odds ratio (used for rank-based enrichment) is derived by performing the Fisher exact test across numerous random gene sets to estimate the mean and standard deviation of the expected rank for each term in the gene-set library. A Z-score is then computed to quantify the deviation of the observed rank from expectation.

### Identification of Disease-Associated Somatic Variants

Disease-associated somatic variant data for *ATXN7L3-AS1* were collected from the COSMIC^94^ and ClinVar^120^ databases via LncBook 2.0^121^. We retained variants labeled as “confirmed somatic mutations” in COSMIC. Variants were classified as pathogenic if they were annotated as having a significant impact on disease in ClinVar or if they had a FATHMM-MKL score greater than 0.7, a threshold previously shown to predict pathogenic potential of noncoding variants^122^ (Table S6A-B).

### Expression of lncRNAs in normal Tissues and TCGA pan-cancer cohorts

The expression profiles of lncRNAs in normal tissues were obtained from the GTEx database, and those in the TCGA Pan-Cancer dataset were obtained through The Atlas of Non-coding RNAs in Cancer (TANRIC)^123^ (Table S7A-D).

### ADDITIONAL RESOURCES

In vivo amplification and characterization of patient-derived xenograft (PDX) tumors have been described in STAR Protocols^59^. Detailed protocols for transduction of PDAC cells and subtype characterization of PDX tumors were published in Heliyon^60^.

### Tissue immunohistochemistry (IHC), cell immunofluorescence (IF) staining, and qRT-PCR

Tissue IHC, IF, and qRT-PCR were performed as previously described in our published work^16^.

## KEY RESOURCES TABLE

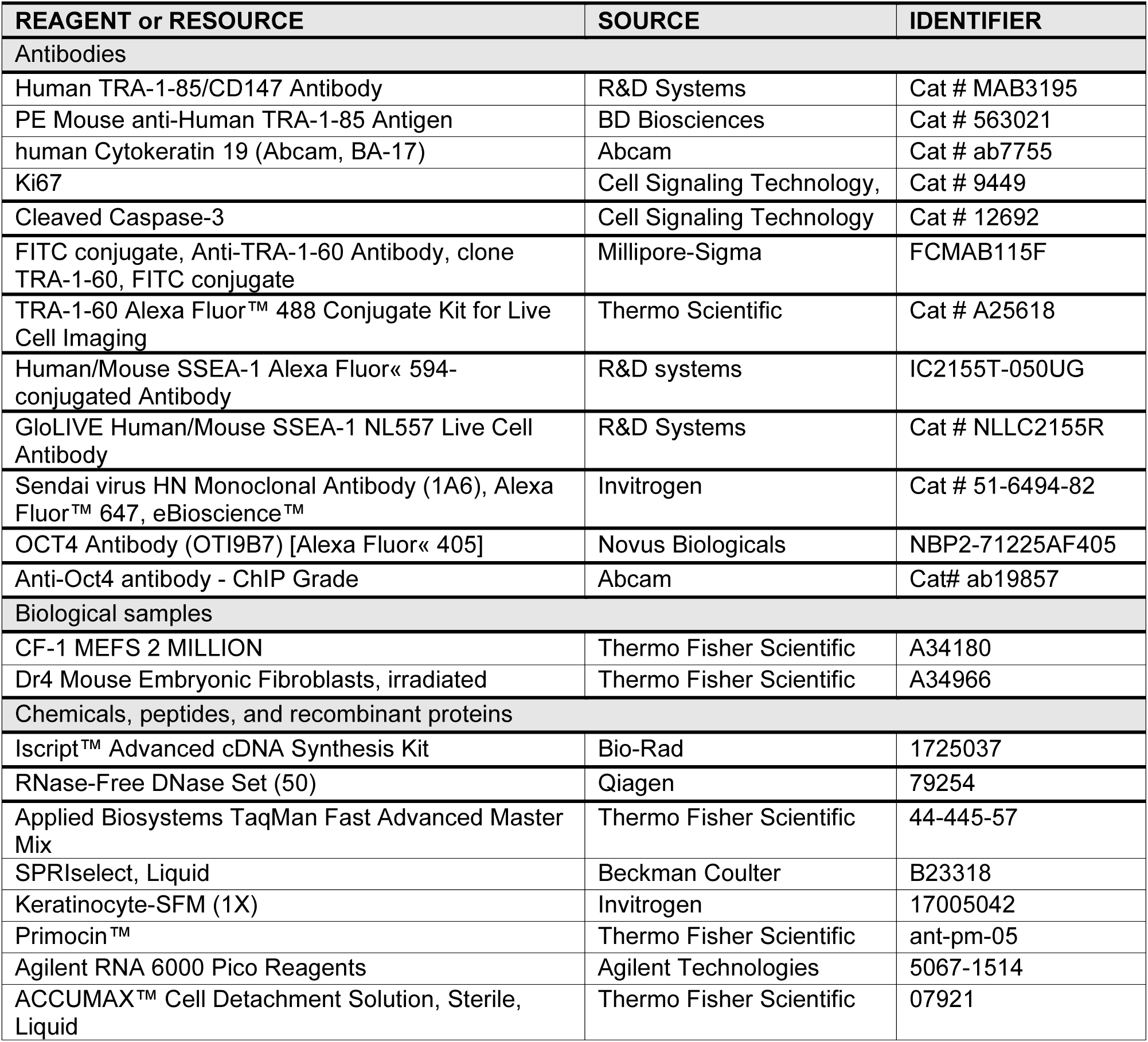
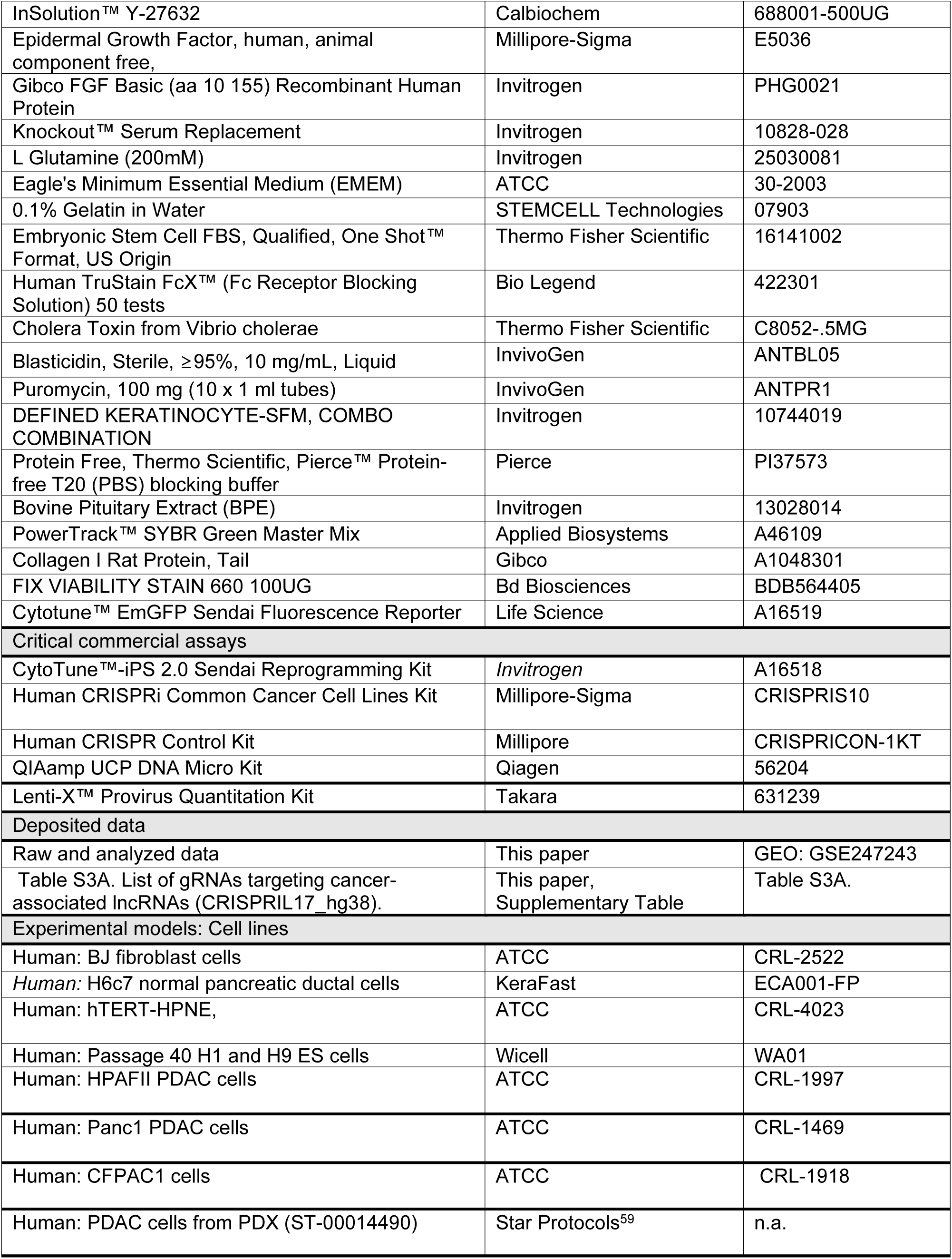
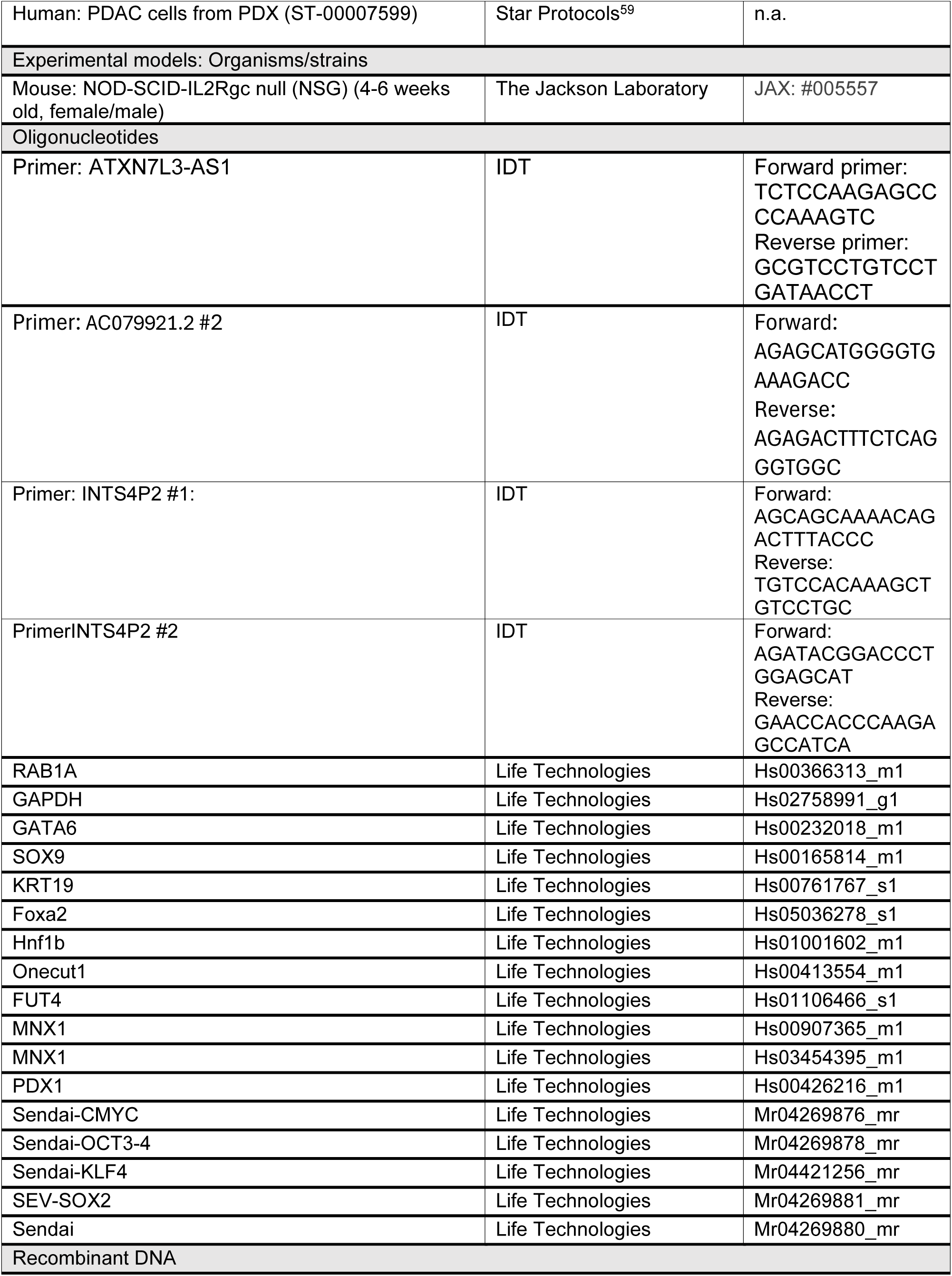
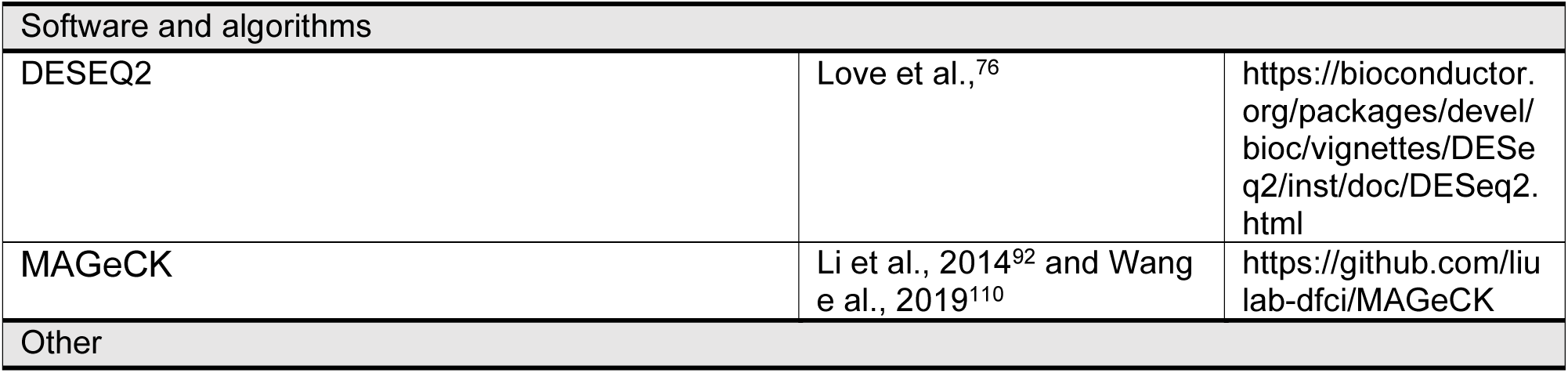

## SUPPLEMENTAL FIGURE TITLES and LEGENDS

**Figure S1(Relevant to Figure 1).**
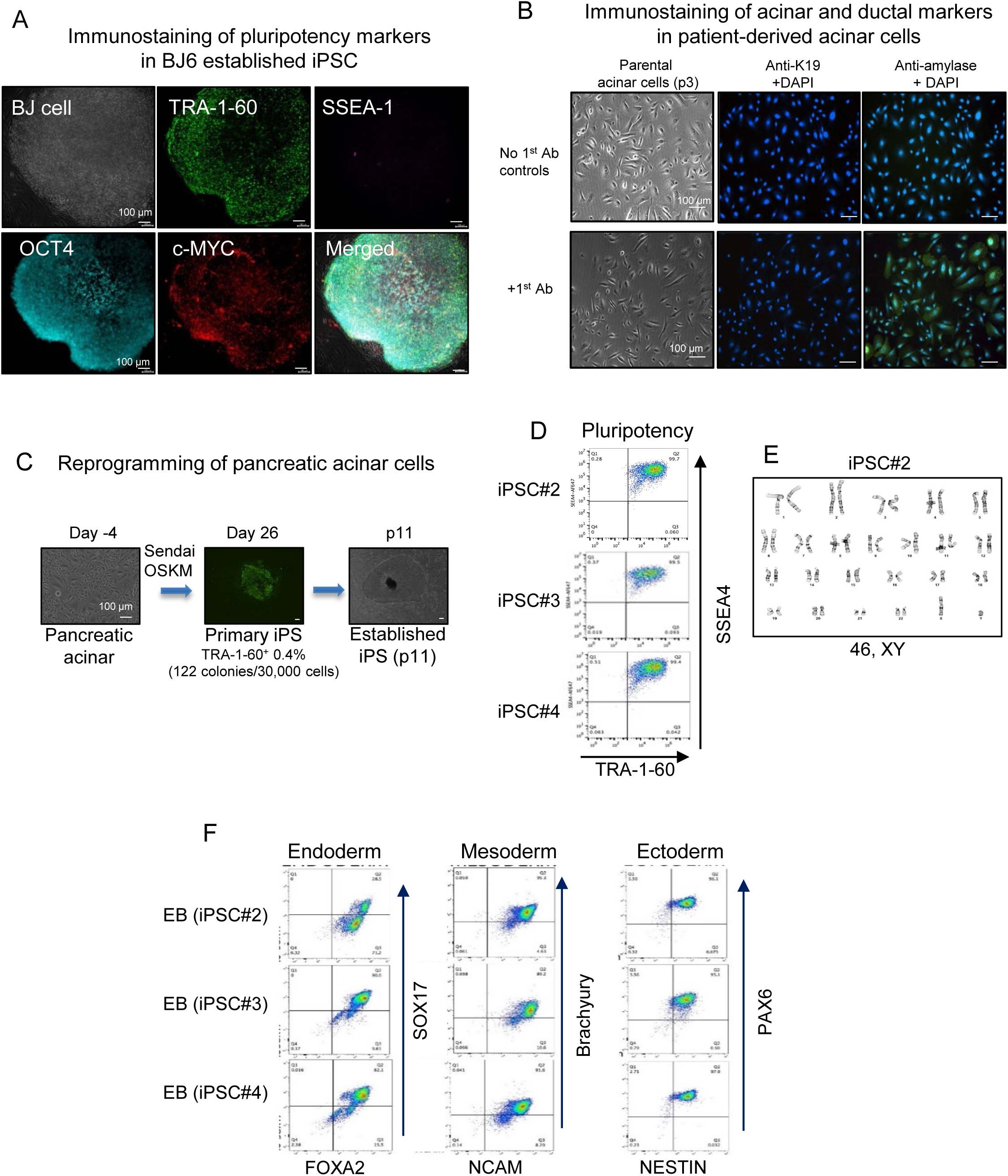
Transgene-free reprogramming of human normal fibroblast and pancreatic acinar cells into pluripotency using Sendai-virus mediated expression of OSKM master TFs. A. Immunostaining of pluripotency markers in iPSCs established from BJ6 human normal fibroblast cells. B. Verification of expression of pancreatic acinar marker amylase and lack of pancreatic ductal marker K19 on pancreatic acinar cells using immunofluorescence (IF). C. Schematic diagram of reprogramming pancreatic acinar cells into iPSC using Sendai-OSKM. Initial numbers of cells and emerged TRA-1-60^+^ cells after 26 days were counted to determine the reprogramming efficiency. Three clones were established after five passages. D. Confirmation of expression of pluripotency surface markers TRA-1-60 and SSEA4 in established acinar iPS lines through flow cytometry. E. Representative image of cytogenetic analysis on 20 G-banded metaphase cells derived from pancreatic acinar-derived iPSC. F. Confirmation of functional pluripotency of acinar-derived iPSC clones. Their embryoid bodies (EB) differentiated into three germ layer lineages, which were verified by staining for three germ layer markers using flow cytometry.

**Figure S2(Relevant to Figure 1).**
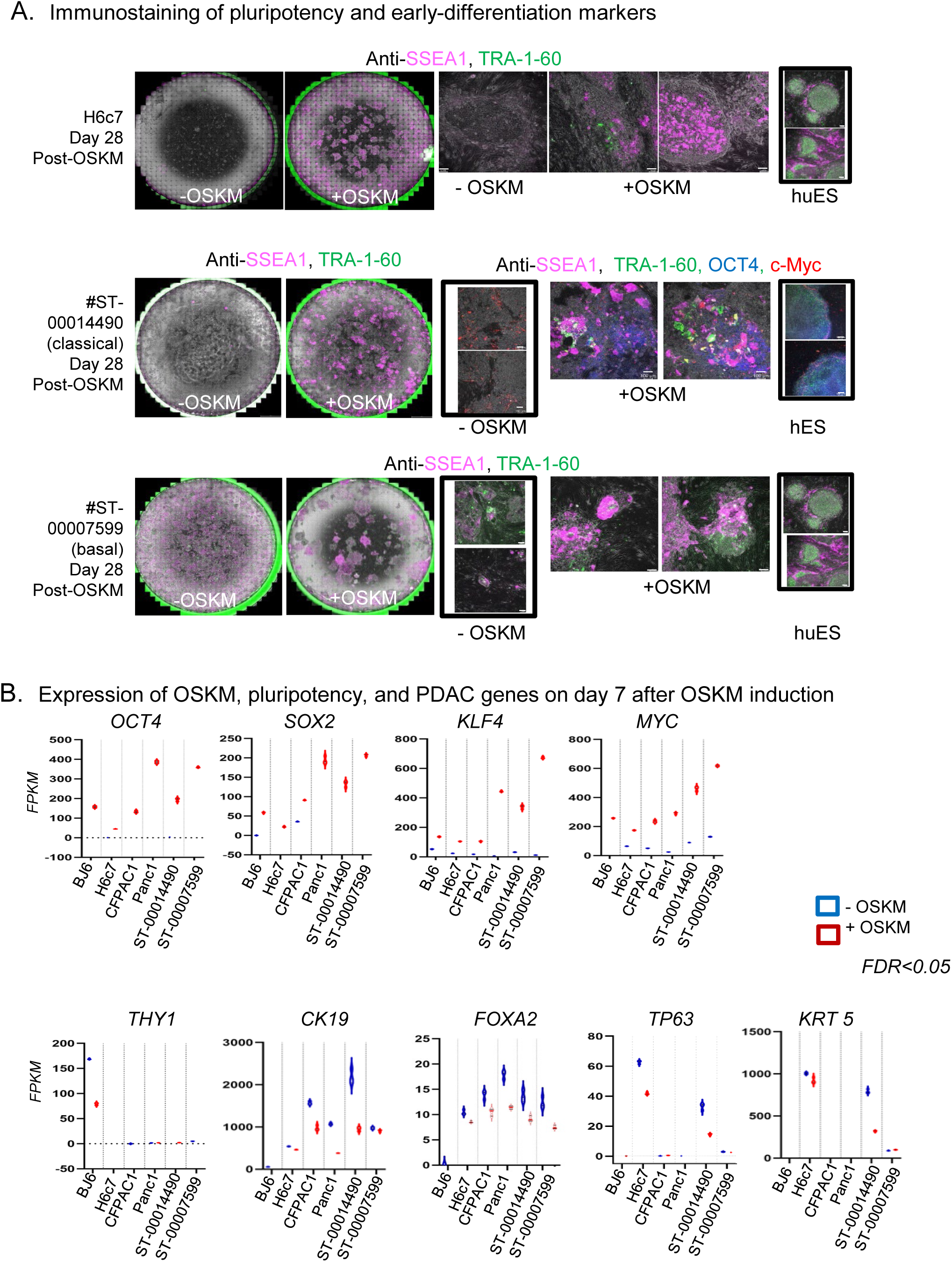
Reprogramming of human normal pancreatic ductal and cancer cells. A. Representative images of colonies stained for TRA-1-60 and SSEA-1 in indicated cells. Scale bars indicate 100 µm. B. Expression of OSKM, pluripotency, and PDAC genes in the indicated cells before and 7 days after OSKM induction.

**Figure S3 (Relevant to Figures 1-2).**
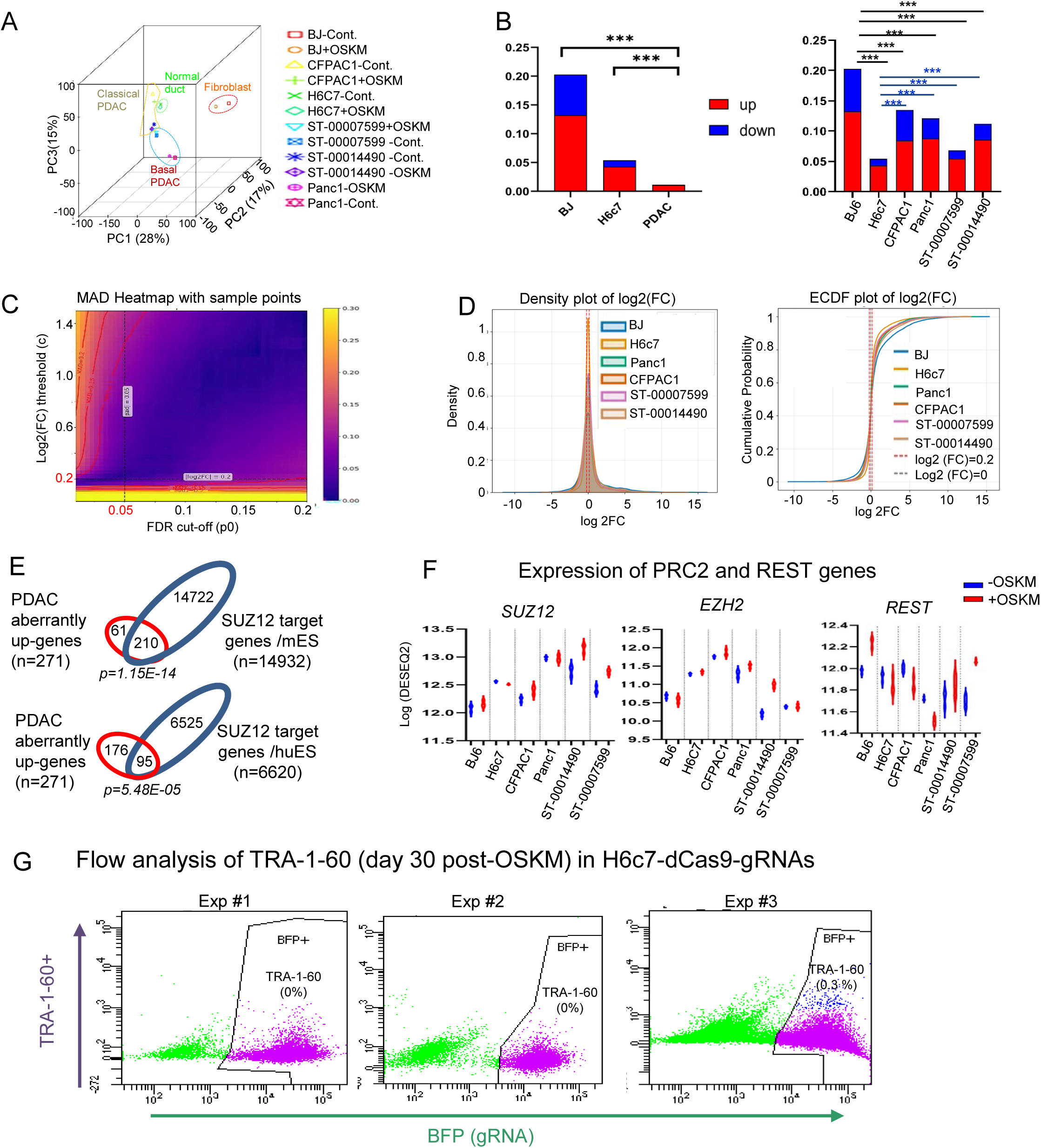
A-F. RNA-seq of classical (CFPAC1 and ST-00014490) and basal (Panc1 and ST-00007599) subtypes of PDAC and normal controls (human BJ6 fibroblast and H6c7 hnPDECs) before and 7 days after OSKM induction. (A) PCA plot. (B) Proportion of genes changed upon OSKM induction among a total of 302,383 genes in normal cells (BJ, H6c7) and PDAC (CFPAC1, ST-00014490, Panc1, and ST-00007599). A chi-squared test was used to compare the proportion of genes exhibiting expression changes following OSKM induction in PDAC cells relative to other normal cell types. The Y-axis indicates the proportion of DEGs relative to the total number of genes. *** indicates p-value < 2.2 × 10⁻^13^. Black and blue bars indicate comparison between PDAC cells and BJ or H6c7 cells, respectively (See Table S2D-E). (C) Mean Absolute Deviation (MAD) heatmap of Q–Q statistics across FDR cut-off points and log₂ (FC) truncated thresholds. The heatmap displays the MAD between observed Q–Q statistics and a truncated standard normal distribution (N(0,1)), computed across combinations of FDR cutoffs (x-axis) and log₂ FC thresholds (y-axis). The color key indicates MAD of Q-Q (stat) vs truncated N (0,1). Lower MAD values (dark blue) indicate better agreement with the expected null distribution, suggesting more reliable gene selection. Red contours mark regions of increasing deviation. Black horizontal and vertical lines indicate log₂(FC) = 0.2 and FDR 0.05 thresholds, respectively. (D) Density and Empirical Cumulative Distribution Function (ECDF) plots of log2 FC values across all samples. Red dashed lines indicate |log2 FC|=0.2. (E) Intersection of aberrantly upregulated PDAC genes (downregulated in normal but upregulated in PDAC) and SUZ12 target genes in human and mouse ES cells (hypergeometric p-value). F. Expressions of genes encoding PRC2 components (SUZ12 and EZH2) and REST transcriptional repressor in indicated cells before and after OSKM induction. G. Flow cytometry analysis of TRA-1-60⁺ cells on day 30 after OSKM induction in H6c7-dCas9 cells expressing gRNAs targeting lncRNAs.

**Figure S4 (Relevant to Figure 3).**
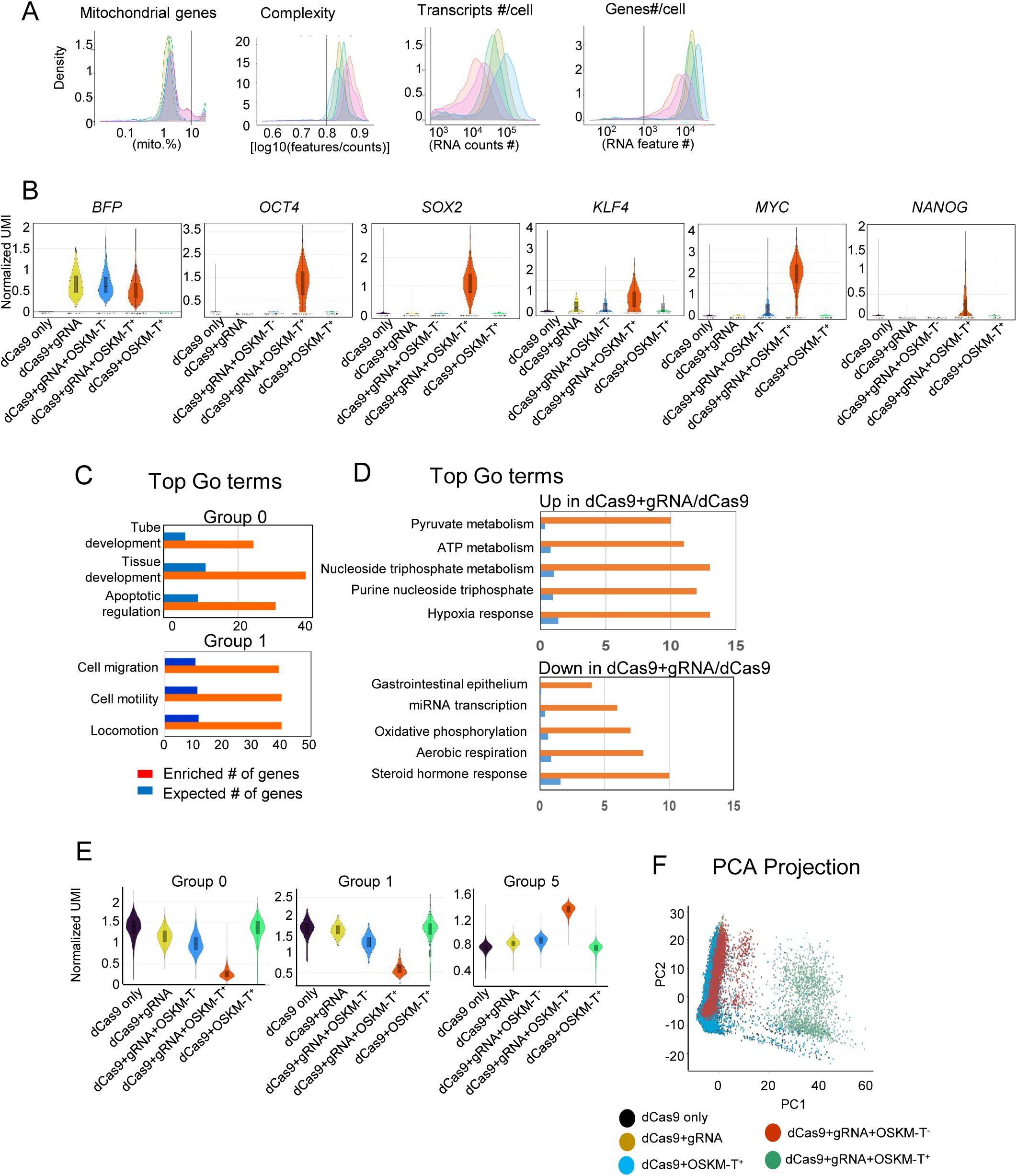
scRNAseq of HPAFII-dCas9+gRNA+OSKM-T^+^ cells along with controls. A. Quality control density plots showing the distribution of cells by key metrics: percentage of mitochondrial gene expression, complexity (features per count), total transcripts per cell, and number of detected genes per cell. B. Expression levels of *BFP*, *OSKM*, and *NANOG* across samples (dCas9 only, dCas9+gRNA, dCas9+gRNA+OSKM-T^−^, dCas9+gRNA+OSKM-T^+^, dCas9+OSKM-T^+^). The X-axis represents individual samples, and the Y-axis shows normalized UMI counts. C. Top GO terms enriched in genes from Group 0 and Group 1 clusters. D. Top GO terms associated with DEGs in dCas9+gRNA cells compared to parental dCas9 cells. E. Violin plots show normalized UMI counts for genes in cells from groups 0, 1, and 5 across samples. F. PCA projection of each sample.

**Figure S5 (Relevant to Figure 4-6).**
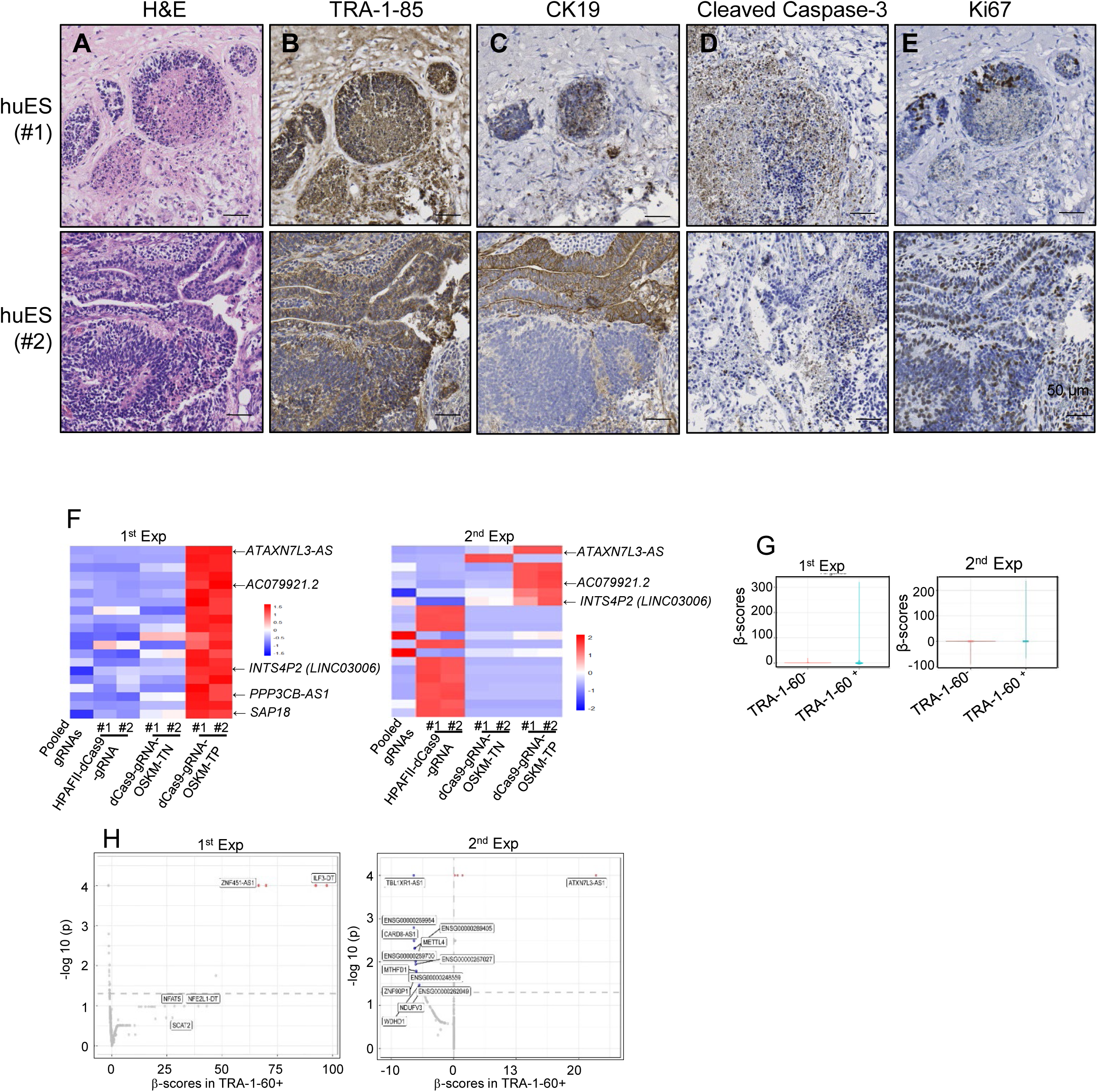
A-E. Xenograft of huES cells; (A) Representative images of immunostaining of H&E, (B) human-specific marker TRA-1-85, (C) human ductal marker CK19, (D) apoptotic marker Cleaved Caspase-3, and (E) proliferation marker Ki67. F-H. Identification of PDAC-enriched lncRNAs depleted during reprogramming; (F) Z-score of top 20 gRNAs. (G) The estimated β-score plots of TRA-1-60^+^ and TRA-1-60^−^ cells compared to references (HPAFII-dCas9-gRNA). The β scores of all gRNAs are normalized based on the negative control normalization. (H) Volcano plot of top-ranked lncRNAs showing β scores enriched in TRA-1-60^+^ cells compared to the reference control. The Y-axis displays –log_10_ (p-values). The p-value is calculated by ranking genes with a skewed distribution compared to a uniform null model and permuting to calculate the significance of the skew in distribution. The X-axis indicates the β scores in TRA-1-60^+^. Red indicates positive enrichment (beta score >0), and blue indicates negative enrichment (beta score <0).

**Figure S6 (Relevant to Figure 6).**
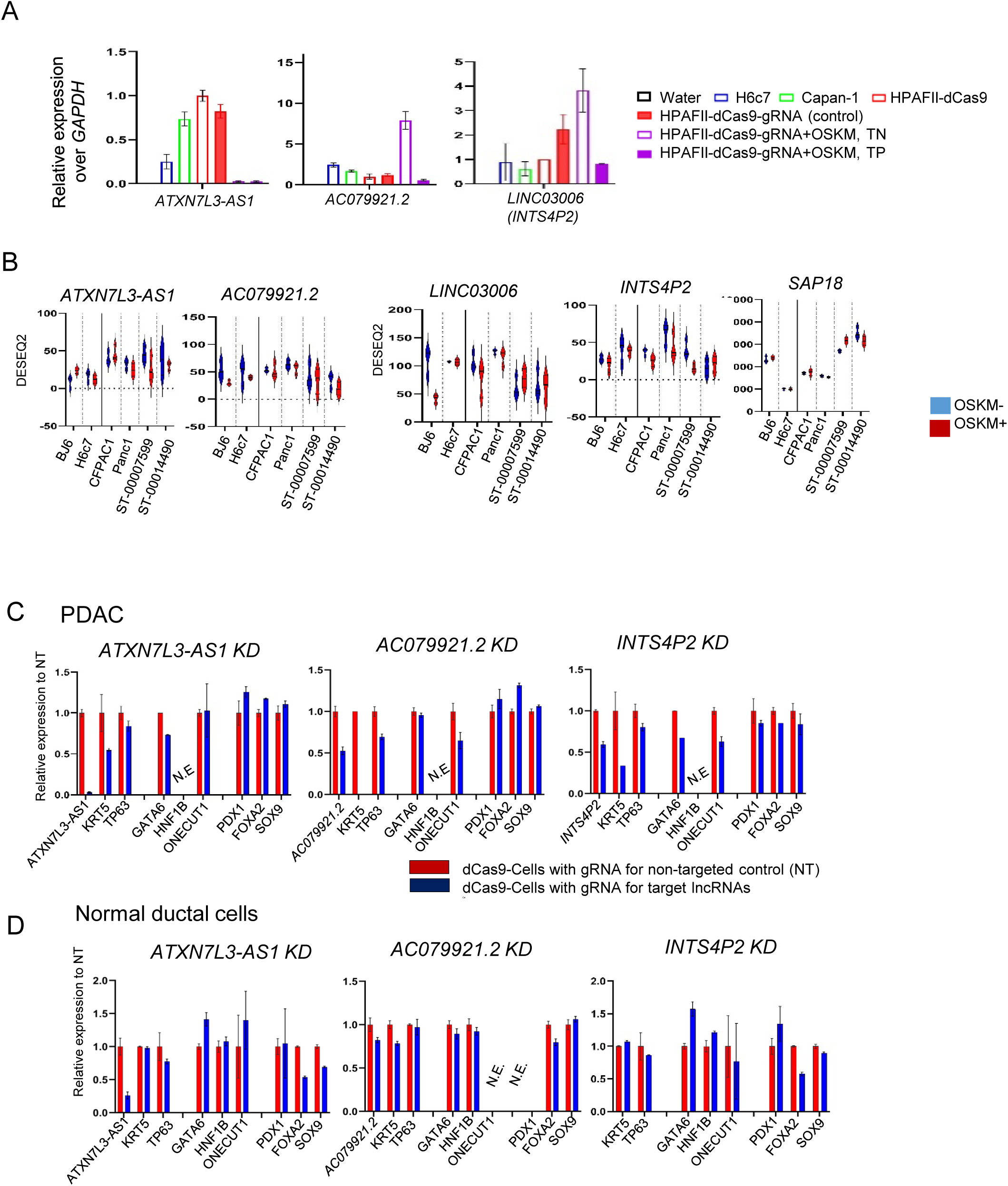
A. The expression of candidate lncRNAs in normal pancreatic cells and cancers, and HPAFII-dCas9+gRNA+OSKM-T^−^ and T^+^ cells (n=3, y-axis indicates relative expression over *GAPDH*, error bars=SD). B. Expression of candidate lncRNAs before and after OSKM induction (n=3, each cell). C-D. Expression of basal subtype markers (*KRT5, TP63*), classical ductal TFs (*GATA6, HNF1B,* and *ONECUT1*), and pancreatic progenitor markers (*PDX1, FOXA2,* and *SOX9*) in PDAC (HPAFII, C) and normal ductal (H6c7, D) cells following CRISPRi-mediated knockdown of *ATXN7L3-AS1*, *AC079921.2*, or *INTS4P2*. Y-axis indicates relative expression normalized to *GAPDH* and scaled to non-targeting controls (NT); error bars represent SD. N=2∼3. N.E. indicates no expression.

**Figure S7 (Relevant to Figure 7).**
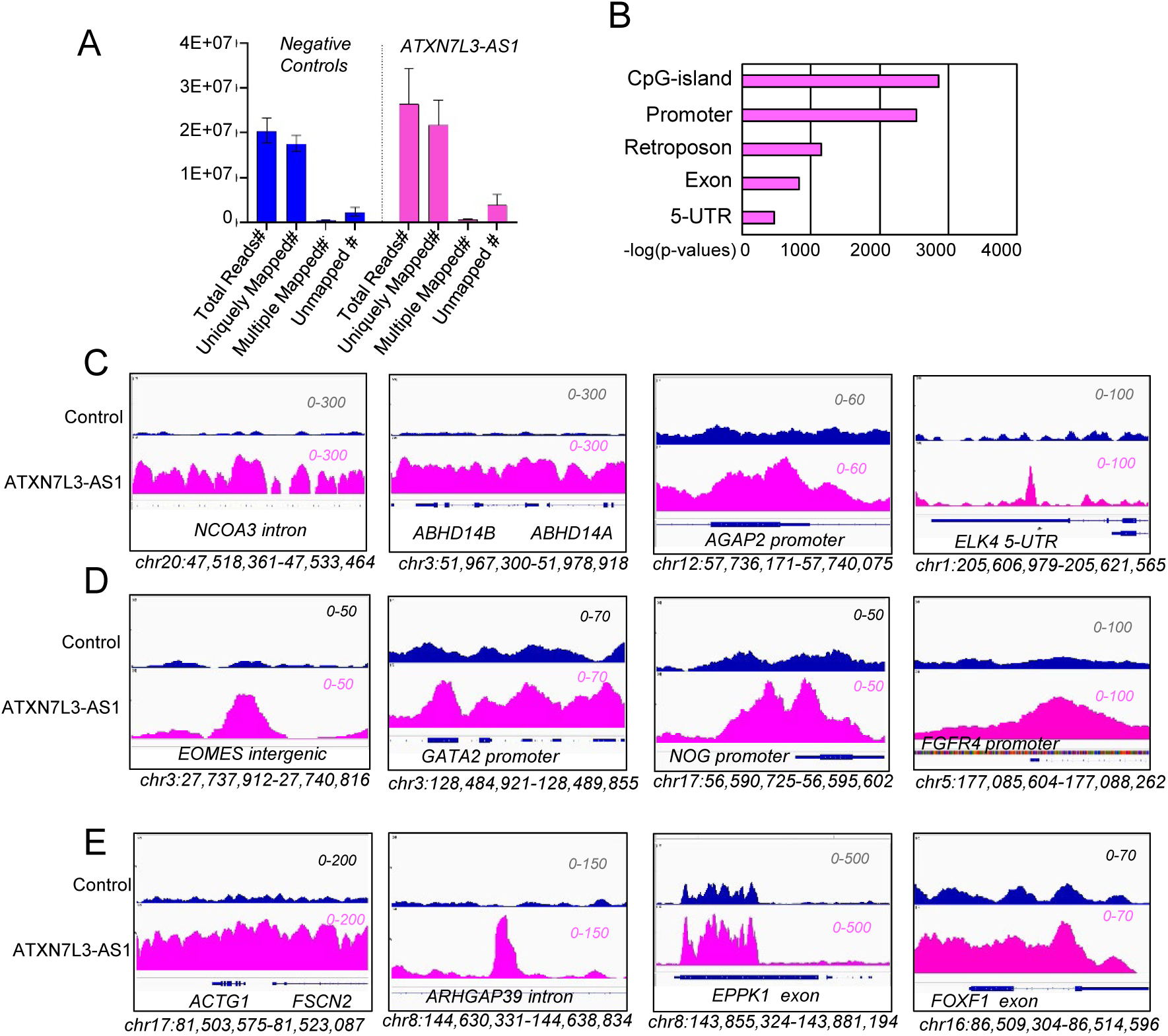
A–E. ChIRP-seq analysis of *ATXN7L3-AS1* genomic occupancy. (A) Read counts from negative control versus pooled *ATXN7L3-AS1* ChIRP samples. (B) Genome-wide distribution of *ATXN7L3-AS1* binding sites. (C–E) Representative genome browser views of *ATXN7L3-AS1* binding over (C) transcriptional activators, (D) mesodermal/neuronal/ ductal developmental genes, and (E) actin cytoskeletal genes.

**Figure S8. (Relevant to Figure 8).**
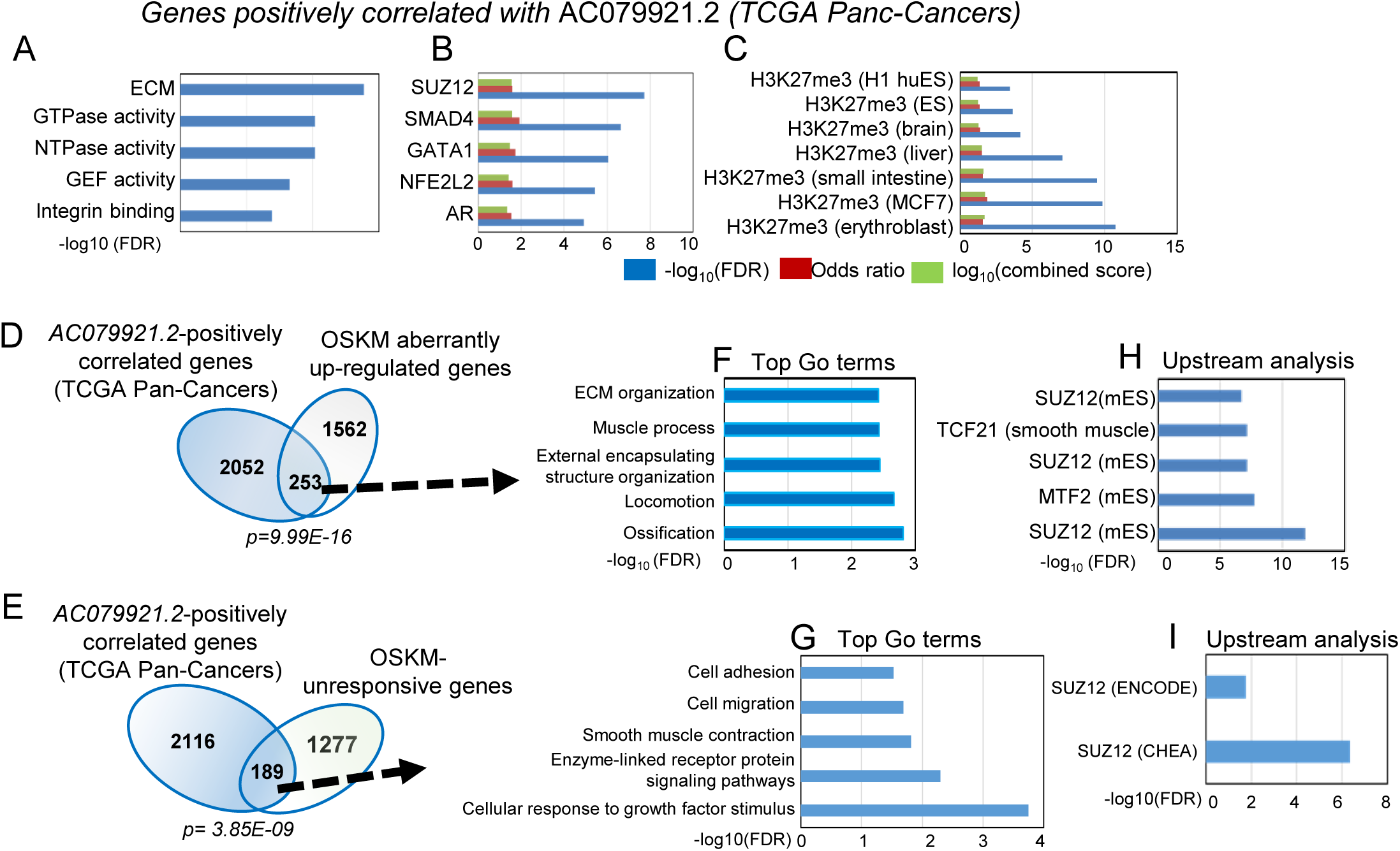
*AC079921.2* Associate with Pro-Invasive Gene Programs. Association of *AC079921.2* with cell adhesion and migration in the TCGA Pan-cancer dataset (TANRIC^107^). A–C Top GO terms (A), potential regulators (B), and histone modifications (C) enriched upstream of genes positively correlated with AC079921.2 in TCGA pan-cancer (Spearman correlation > 0.4; n = 2,305, Table S7B). D-E. Overlap of *AC079921.2*–positively correlated genes in pan-TCGA tumors with OSKM-aberrantly upregulated (D) or unresponsive (E) genes in PDAC reprogramming. The hypergeometric test was used for significance. F-G. GO analysis of overlapping genes from (D) and (E), enriched for ECM organization, muscle contraction, cell adhesion, and migration pathways. H-I. Upstream regulator analysis identifies SUZ12 and chromatin-associated complexes as potential regulators of AC079921.2–associated transcriptional programs.

## SUPPLEMENTAL TABLE TITLES and LEGENDS

**Table S1.** (*Related to Figure 1*) Table S1. KRAS genotyping of iPSCs derived directly from primary PDAC samples.

**Table S2.** (*Related to Figure 1*) OSKM-mediated reprogramming is not sufficient to override PDAC identity.

- **Table S2A.** Regularized log₂-transformed DESeq2 expression values for 375 PDAC-unique genes.
- **Table S2B.** Regularized log₂-transformed DESeq2 expression values for 7 PDAC-unique genes regulated by OSKM induction.
- **Table S2C.** Gene ontology (GO) analysis of 7 PDAC-unique genes downregulated by OSKM induction.
- **Table S2D.** Chi-square test comparing the proportion of genes with expression changes in pooled PDAC vs. normal cells following OSKM induction.
- **Table S2E.** Chi-square test comparing the proportion of genes with expression changes in each PDAC and normal sample following OSKM induction.
- **Table S2F.** List of OSKM-unresponsive PDAC genes (signature: [−1][−1][0][0][0][0], outliers removed, n=1,466, distance < 0.375; related to Fig. 1F).
- **Table S2G.** List of OSKM-aberrantly upregulated PDAC genes (signature: [−1][− 1][0/+1][0/+1][0/+1][0/+1], outliers removed, n=1,815, distance < 0.25; related to Fig. 1F).
- **Table S2H.** List of strongly OSKM-aberrantly upregulated PDAC genes (signature: [− 1/0][−1/0][1][1][1][1], n=82, distance = 0; related to Fig. 1F).

**Table S3.** (*Related to Figure 2*) Depletion of cancer-associated lncRNAs enhances reprogramming of PDAC.

- **Table S3A.** List of gRNAs targeting cancer-associated lncRNAs (CRISPRIL17_hg38).

**Table S4.** (*Related to Figure 3)* Single-cell RNA-seq reveals transcriptional reprogramming toward pluripotency, suppression of PDAC identity, and reactivation of tumor suppressor genes (TSGs)

- **Table S4A.** List of pluripotency marker genes.
- **Table S4B.** List of endodermal and PDAC marker genes.
- **Table S4C.** List of ectodermal and mesodermal marker genes.
- **Table S4D.** List of “PDAC-specific signature genes” (n=332)
- **Table S4E**. List of OncoKB-annotated tumor suppressor genes (total: n=379; excluding oncogenes: n=318).

**Table S5.** (*Related to Figure 6)* Identification of depleted gRNAs in HPAFII-dCas9+gRNA+OSKM-T^+^ cells. sample

- **Table S5A.** Sample barcodes and Illumina adaptor sequences.
- **Table S5B.** Read counts aligned to gRNAs (Experiment 1).
- **Table S5C.** Read counts aligned to gRNAs (Experiment 2).
- **Table S5D.** Raw hit counts (Experiment 1).
- **Table S5E.** Raw hit counts (Experiment 2).
- **Table S5F.** Z-scores of each gRNA (Experiment 1).
- **Table S5G.** Z-scores of each gRNA (Experiment 2).
- **Table S5H.** MAGeCK-MLE output (aggregated reference x500, Experiment 1).
- **Table S5I.** MAGeCK-MLE output (Experiment 2).
- **Table S5J.** MAGeCK-MLE output (combined Experiment 1 and 2).
- **Table S5K.** Final list of selected candidate gRNAs.

**Table S6.** *(Related to Figure 6-7)* COSMIC Pathogenic somatic variants in *ATXN7L3-AS1* (GRCh38) and ChIRP-Seq for *ATXN7L3-AS1*.

- **Table S6A**. Pathogenic somatic variants in *ATXN7L3-AS1* identified in pan-cancers.
- **Table S6B**. Pathogenic somatic variants in *ATXN7L3-AS1* identified in pancreatic cancer.
- **Table S6C**. *ATXN7L3-AS1* CHIRP probes
- **Table S6D**. *ATXN7L3-AS1* ChIRP-Seq DNA Mapping
- **Table S6E**. Normalized DESeq2 counts of genes in Group 1 (red; basal subtype– associated, n = 552).
- **Table S6F**. Normalized DESeq2 counts of genes in Group 2 (yellow; normal pancreatic ductal cell–associated, n = 460).
- **Table S6G**. Normalized DESeq2 counts of genes in Group 3 (blue; fibroblast-associated, n = 702). sample
- **Table S6H**. Normalized DESeq2 counts of genes in Group 4 (purple; cancer-associated, n = 1,839).

**Table S7**. (Related to Figures 7 and S8) TCGA Pan-Cancer Genes Correlated with *ATXN7L3-AS1* and *AC079921.2*.

- **Table S7A.** TCGA Pan-Cancer Genes Positively Correlated with *ATXN7L3-AS1* (Spearman correlation > 0.4)
- **Table S7B.** TCGA Pan-Cancer Genes Positively Correlated with *AC079921.2* (Spearman correlation > 0.4)
- **Table S7C.** TCGA Pan-Cancer Genes Negatively Correlated with *ATXN7L3-AS1* (Spearman correlation <-0.4)
- **Table S7D.** TCGA Pan-Cancer Genes Negatively Correlated with *AC079921.2* (Spearman correlation <-0.4)

## Notes

### Competing Interest Statement

The authors have declared no competing interest.

### Summary of Updates

Data and abstract have been updated.

